# CtIP-dependent nascent RNA expression flanking DNA breaks guides the choice of DNA repair pathway

**DOI:** 10.1101/2022.01.14.476291

**Authors:** Daniel Gómez-Cabello, Giorgios Pappas, Diana Aguilar-Morante, Christoffel Dinant, Jiri Bartek

**Affiliations:** Genome Integrity Unit, Danish Cancer Society Research Center, Strandboulevarden 49, Copenhagen, DK-2100, Denmark; Instituto de Biomedicina de Sevilla (IBiS), Hospital Universitario Virgen del Rocío/CSIC/Universidad de Sevilla, 41013 Seville, Spain; Departamento de Genética, Facultad de Biología, Universidad de Sevilla, 41012 Seville, Spain; Division of Translational Medicine and Chemical Biology, Department of Medical Biochemistry and Biophysics, Science for Life Laboratory, Karolinska Institute, Scheele’s vag 2, Stockholm, 17177, Sweden

**Keywords:** DNA repair, Homologous recombination, DNA damage, Double-Strand Break, Transcription, RNA polymerase II inhibitor, DNA resection, cancer

## Abstract

The RNA world is changing our views about sensing and resolution of DNA damage. Here, we developed single-molecule DNA/RNA analysis approaches to visualize how nascent RNA facilitates the repair of DNA double-strand breaks (DSBs). RNA polymerase II (RNAPII) is crucial for DSB resolution in human cells. DSB-flanking, RNAPII-generated nascent RNA forms RNA:DNA hybrids, guiding the upstream DNA repair steps towards favouring the error-free Homologous Recombination (HR) pathway over Non-Homologous End Joining. Specific RNAPII inhibitor, THZ1, impairs recruitment of essential HR proteins to DSBs, implicating nascent RNA in DNA end resection, initiation and execution of HR repair. We further propose that resection factor CtIP interacts with and re-activates RNAPII when paused by the RNA:DNA hybrids, collectively promoting faithful repair of chromosome breaks to maintain genomic integrity.

## Introduction

The concept of an RNA world postulates that RNA was essential for molecular processes and biochemical reactions implicated in the origin of life on Earth (1). To compensate for RNA instability, DNA appeared later during the evolution to better preserve genetic information, followed by fidelity mechanisms to maintain genome stability (1). Recently, RNA has emerged as a major factor in essential mechanisms regulating gene expression (2,3) and contributing actively to DNA repair processes (4-9). Arguably the most cytotoxic genomic lesions are DNA double-strand breaks (DSBs), lesions repaired mainly by either of the two major pathways: non-homologous end-joining (NHEJ) and homologous recombination (HR) (10-14). While numerous protein components of these two pathways have been discovered over time, only recently RNA has been implicated in DSB repair as well. For example, recent evidence showed that DSBs in transcriptionally active genomic regions are more prone to be repaired by HR (7,8,15).

Interestingly, only 2-8% of the human genome ever get transcribed (16), yet it is unclear, unlikely perhaps, that HR is restricted to these genomic regions only. Hence, chromatin structure at transcriptionally active sites and the influence of diverse, relevant mechanisms, including DNA repair pathways, are currently subject to intense investigation to elucidate to what extent and how RNA impacts DSBs repair. So far, such efforts generated controversial results. On the one hand, global RNA transcription is inhibited after DNA damage to avoid conflicts between repair and other DNA metabolic processes such as replication (17). RNA in DNA repair processes has also been demonstrated to contribute to genomic instability by forming RNA:DNA hybrid structures (4,18).

On the other hand, the formation of RNA:DNA hybrids regulates DNA repair in diverse organisms (7,8,19), exemplified by the DNA damage response RNAs (DDRNAs), necessary for DDR activation (20-22). Furthermore, recruitment of general transcription factors to DNA damage sites was reported, and an active role for RNA polymerase II (RNAPII) in DNA repair was proposed (23). Recently, RNA polymerase III was reported to be actively recruited to DSBs by the MRN complex and mediates RNA synthesis, promoting HR repair (24). However, any role of RNAPII, the major mammalian RNA polymerase, in this context remains unknown. Any potential mechanistic contribution to DNA repair pathway choice could also inspire new cancer treatment strategies to be combined with standard-of-care DNA damaging radio-chemotherapy.

As the repair mode is critical for genomic integrity and thereby inheritance, evolution, organismal development and tissue homeostasis, the emerging evidence for RNA involvement raises the crucial questions of whether and how could RNA guide the choice between HR and NHEJ in DSB repair, an issue that we address in our present study.

Here, we elucidate the role played by *de novo* RNA synthesis by RNAPII in DSB repair *via* HR in human cells. We found that RNA presence during different cell cycle phases impacts the decision between the HR and NHEJ repair pathways, combined with the homologous sequences of sister chromatids. Using our innovative single-molecule analysis approaches, employed here for the first time, we observed nascent RNA overlapping with ssDNA resection tracts generated during DNA end resection, indicating that RNAs, mainly synthesized by RNAPII, are essential to initiate DNA end resection and thereby shift the choice of DSB repair towards using the more faithful HR over NHEJ. Indeed, RNA:DNA hybrid formation is essential for resection processing and as a repair regulatory step in the HR pathway. Moreover, RNAPII inhibition, using a specific CDK7 inhibitor THZ1, impairs HR factor recruitment to DSB. We further demonstrate a previously unsuspected function of CtIP and BRCA1 as transcription re-activators of RNAPII paused transiently by the RNA:DNA hybrids during the early-stage response to DSBs, thereby promoting DNA resection and skewing DSB repair balance towards HR.

## Material and Methods

### Cell culture

U2Os and Hela cells were cultured in high-glucose Dulbecco’s Modified Eagle Medium (DMEM) plus GlutaMax supplemented with 10% FBS, 100 μg/ml streptomycin and 100 U/ml penicillin at 37°C in 5% CO2. siRNAs against CtIP (GCUAAAACAGGAACGAAUC), BRCA1 (GGAACCUGUCTCCACAAAG), RAD52 (s11747, ThermoFisher) and a control Non-Target sequence (Sigma Aldrich) were transfected with RNAiMax lipofectamine reagent mix (Life Technologies), according to the manufacturer’s instructions.

### Immunofluorescence

U2OS cells were grown on coverslips and pre-extracted for 5 min on ice using 0.2% Triton X-100 in PBS, then fixed with 4% paraformaldehyde (w/v) in PBS for 15 min, washed three times with PBS and blocked for at least 1 hour with 5% FBS diluted in PBS. Cells were incubated with the adequate primary antibodies (Supp. Material 1), diluted in 5% FBS in PBS for 16h at 4°C, washed with PBS, and then incubated with secondary antibodies (Supp. Material1) diluted in 5% FBS in PBS for 1 hour at room temperature (RT). Cells were then washed twice with PBS, and coverslips were mounted with Vectashield mounting medium (Vector Laboratories) containing 4’,6-diamidino-2-phenylindole and analyzed using a LEICA microscope. At least 2000 cells were scored per sample. Experiments were repeated at least three times independently.

### High Content image acquisition

Quantitative image-based cytometry (QIBC) was performed as previously described (25). Images were acquired in an automatic and unbiased way by the scanR acquisition software and analysed by scanR image analysis software. Results were exported as txt files. The txt dataset was further processed with spotfire and PRISM software (Graphpad Software Inc) for further analysis. Statistical significance was determined with an Ordinary one-way and two-ways ANOVA tests using multiple comparison by PRISM software (Graphpad Software Inc). Statistically significant differences were labelled with one, two or three asterisks if *p*< 0.05, *p*< 0.01 or *p* < 0.001.

### Metabolic labelling of nascent RNA by EU

Nascent RNA was visualised using metabolic labelling using Invitrogen™ Click-iT™ RNA Alexa Fluor™ 594 Imaging Kit, with modifications. Briefly, cells were cultured in complete media and pulsed for 30 min with EU at a final concentration of 1 mM, before fixation with 4% paraformaldehyde (PFA) for 10 min. After fixation, cells were permeabilised with 0.5% Triton X-100 and washed 3 times with Tris-buffered saline (TBS) (50 mM Tris pH 8.0, 150 mM NaCl). Click reaction master mix was then prepared as follows: 5 µM Alexa Fluor 488 azide, 2 mM CuSO4, 100 mM Sodium ascorbate before samples preparation for acquisition using QIBC. Nucleoli EU mean intensity was analyzed using spot detector tool, and was excluded from nucleus EU intensity for determination of Nucleoplasm EU intensity.

### R-SMART

Hela and Hela-RNAseH1 cells downregulated for the indicated genes were grown in the presence of 10 μM bromodeoxyuridine (BrdU, GE Healthcare) for 24 hours. Cultures were then irradiated (5 Gy) and incubated with 100 μM RNA precursor 5-ethynyluridine (EU) and harvested after 1 hour. Cells were lysed using Spreading Buffer (200 mM Tris:HCl pH 7.5, 50 mM EDTA, 0,5% SDS). 2000 cells were used to stretch nucleic acid fibres on coverslips using a 15° angle of incline as previously described. Five slides were stretched for all experimental conditions, and two or three slides for each condition were stained. Nascent RNA was detected using Click-it RNA Alexa Fluor 488 Imaging Kit (ThermoFisher Scientific) following the manufacturer’s instructions. Then samples were incubated directly without denaturation with an anti-BrdU mouse monoclonal antibody (Becton Dickinson, 347580). Secondary antibodies were DayLight 555 anti-mouse (Thermo Fisher Scientific).

Images of DNA fibres were acquired using an LSM800 confocal microscope (Carl Zeiss) and a Plan-Apochromat 63×/1.4 numerical aperture (NA) oil immersion objective (Carl Zeiss). Labelled RNA and DNA fibres were analyzed using LSM ZEN software, using a colocalization tool to measure pixels staining for both fluorescence. More than 20 fields or more than 100 fibres were scored during measurements on each slide, for every repeated experiment. The percentage of RNA and DNA resection staining pixels and their colocalization were presented from both experiments.

### RL-SMART

Hela and Hela-RNAseH1 cells were treated with 10 μM bromodeoxyuridine (BrdU, GE Healthcare) for 24 hours. Cultures were then irradiated (5 Gy) harvested after 1 h. Cells were lysed using Spreading Buffer (200 mM Tris:HCl pH 7.5, 50 mM EDTA, 0,5% SDS). 2000 cells were used to stretch nucleic acid fibres as described in the R-SMART section. Then samples were incubated directly without denaturation with an anti-BrdU rat monoclonal antibody (Becton Dickinson, 347580) and S9.6 mouse monoclonal antibody (Kerafast, ENH001). Secondary antibodies were DayLight 550 anti-rat and 488 anti-mouse. Five slides were stretched for all experimental conditions, and two or three slides for each condition were stained. Images were taken as described above for the R-SMART technique. The percentage of pixels of RNA:DNA hybrids and DNA resection staining, and their colocalization were analyzed as in the R-SMART technique and presented from both experiments.

### DNA Damage by laser micro-irradiation

Cells were plated on 1 glass-bottom well chamber one day before the analysis. Before laser irradiation, cells were incubated with 1 µM THZ1 (1604810-83-4) for 1 hour, at 37°C and 5% CO2. Then, cells were maintained under the same conditions using Temperature Control Chamber (PerkinElmer UltraView VoX), and images were taken using Nikon Eclipse Ti microscope equipped with a 63x oil objective and collected every 4 seconds for 10 minutes. GFP signal intensity was measured using ImageJ software for at least 20 cells per condition from three biological replicates.

### *In situ* labelling of newly-synthesized RNA with 5’-Bromouridine 5’-Triphosphate

An appropriate number of cells was seeded on coverslips to reach 75% of confluency on the day of the experiment. Coverslips were washed once with PBS and incubated with permeabilization buffer (20 mM Tris-HCl, pH 7.4, 5 mM MgCl2, 0.5 mM EGTA, 25% glycerol, 0.05% Triton X-100, 1 mM PMSF and ribonuclease inhibitor 20 U/ml) for 2 minutes at RT. Next, the permeabilization buffer was removed from the coverslips and transcription buffer (20 mM Tris-HCl, pH 7.4, 5 mM MgCl2, 0.5 mM EGTA, 25% glycerol, 1 mM PMSF, 100 mM KCl, ribonuclease inhibitor 20 U/ml, 500 μM BrUTP, 500 μM CTP, 500 μM GTP and 2 mM ATP) was added. Coverslips were incubated for 8-10 minutes at 37°C. After removing the transcription buffer, coverslips were gently washed with cold PBS and fixed in 4% paraformaldehyde for 10 minutes at RT. Indirect immunofluorescence using mouse anti-BrdU antibody (BD347580) was carried out to detect the incorporation of BrUTP analogue in nascent transcripts.

### Immunoblotting

Whole extracts from cell lines were prepared in Laemmli buffer (4% SDS, 20% glycerol and 125 mM Tris-HCl, pH 6.8), and proteins were resolved using SDS-PAGE and transferred to nitrocellulose membranes, followed by immunoblotting. Western blot analysis was carried out using the antibodies listed in Supp. Material 1. Results were acquired using the ChemiDoc system and visualized with Image Lab Software (Bio-Rad).

### Co-immunoprecipitation assay

U2OS cells were X-irradiated and incubated for specific time points. Nuclear protein fractions were prepared by using Nuclear Complex co-IP kit (Active Motif #5400), according to the manufacturer. Nuclear extracts were subsequently diluted in appropriate volume of IP buffer (0.5% NP40, 50mM Tris-HCl, pH 7.4, 150mM NaCl, 1mM EDTA) supplemented with protease and phosphatase inhibitors. Diluted nuclear extracts were incubated with anti-RNA polymerase II CTD repeat YSPTSPS antibody (8WG16)-ChIP grade (Abcam, ab817) overnight at 4°C. Protein extracts were then incubated with 25 μl of protein G dynabeads (Thermo Fisher, 10004D) for 1 hour at 4°C. Beads were washed 5 times with 250 μl of IP buffer, and 20 μl of Laemmli sample buffer was added combined with heating at 97°C for 5 minutes to elute the precipitated protein fraction from the beads.

## Results

### HR factors impact ionizing radiation-induced nascent RNA

To study the relationship between nascent RNA and DNA repair, we analyzed nascent RNA expression in different cell cycle phases using a modified nucleotide, 5-ethynyl uridine (EU) (26), in human U2OS osteosarcoma cells. We focused on the initial minutes immediately after irradiation as we believe that is the period critical for the cell’s choice to repair DSBs via either NHEJ or HR. Hence our experiments were designed to analyze particularly the initial 30 minutes post-radiation exposure (Fig.1a). In control, proliferating and non-irradiated cells, global RNA transcription in the nucleoplasm increased during the cell cycle, reaching maximum levels in S and G2 phases in undamaged cells (Fig.1b). The overall pattern of nascent RNA transcription was similar in irradiated cells, yet with a significant increase of nascent RNA expression occurring in each and every cell cycle phase, compared to non-irradiated controls (Fig.1a-c). Such RNA increase was transient, returning to pre-irradiation levels by 60 minutes after radiation exposure (Supp. Fig. 1a). HR repair is more active in S and G2 phases due to availability of sister chromatids, yet we wondered whether DNA resection, as a critical upstream step in HR, follows a similar pattern related to nascent RNA transcription. Detection of ssDNA using BrdU detection under non-denaturing conditions by immunofluorescence showed that following irradiation with 5Gy, DNA end resection occurs mainly in early and late S phases, decreasing robustly in G1 and G2, thereby indicating similar patterns of DNA end resection tracts and nascent RNA expression (Fig. 1c-d), and raising the possibility of a functional link between these two processes during cell cycle progression. The presence of CtIP and BRCA1 proteins promote DNA end resection at DSBs (27). To determine any potential role of these proteins in *de novo* RNA transcripts formation after DNA damage, we examined nascent RNA in CtIP-and BRCA1-depleted cells versus mock-depleted controls, under both non-irradiated and irradiated conditions (Fig. 1e and Supp. Fig.1b-d). We used previously published, validated siRNA sequences against these two genes (26-28) for these experiments. Whereas CtIP- and BRCA1-depleted non-irradiated cells showed unaltered global RNA transcription at 30 min and 60 min (Fig. 1b and Supp. Fig. 1b), the irradiated CtIP- and BRCA1-depleted cells showed a significant decrease of nascent RNA expression associated with IR-induced DNA lesions (Fig. 1e,f).

**Fig. 1.**
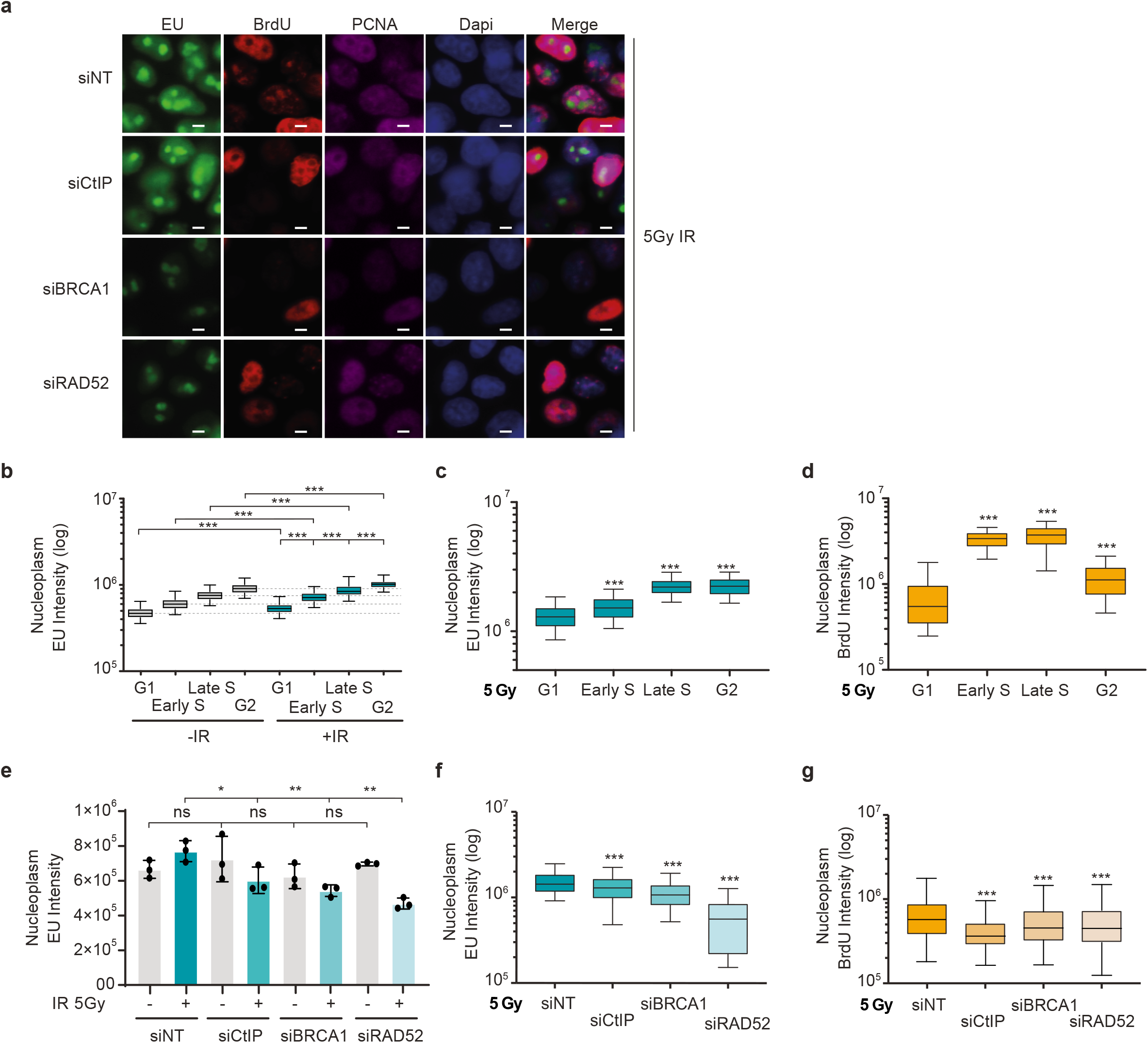
DNA resection (ssDNA) correlates with nascent RNA synthesis. **a** Representative immunofluorescence images of EU, BrdU, PCNA and DAPI staining in U2OS cells depleted for the indicated DDR factors, irradiated with 5Gy and stained upon 30-minute labelling with EU and BrdU that started immediately after IR exposure. Scale bar: 10μm. **b** Graph shows nucleoplasm EU intensity in different cell cycle phases in non- and irradiated-U2OS cells. EU component was added upon DNA damage using 5Gy in irradiated cells and labelled for 30 minutes in both cellular conditions. **c** Graph shows EU intensity in different cell cycle phases in control U2OS cells irradiated and labelled for 30 min as in (a). **d** Graph shows intensity of BrdU staining under non-denaturing conditions to visualize stretches of ssDNA, as a DNA resection marker in cell cycle phases of U2OS cells irradiated. **e** Bar graph represents nucleoplasm EU intensity in siRNAs against indicated genes in non-and irradiated-cells. EU component was added upon DNA damage using 5Gy in irradiated cells and labelled for 30 minutes in both cellular conditions. Showing mean values from 3 independent experiments. Error bars represent s.e.m from at least 500 cells. ns=non-significant, *p>0.05, **p>0.01 and *** p>0.001 using multiple comparison with Ordinary Two-Ways ANOVA. **f** Nascent RNA synthesis in U2OS cells depleted for the indicated DDR proteins, labelled with EU for 30 minutes starting after irradiation with 5 Gy. **g** Quantification of BrdU staining under non-denaturing conditions to mark ssDNA as a DNA resection marker in U2OS cells depleted for CtIP, BRCA1 and RAD52, respectively, and stained after 30-min BrdU labeling started after exposure to 5 Gy. Statistical data at b-d and f-g showing data from 3 independent experiments. Error bars represent Max and Min values from at least 500 cells. ns=non-significant, *p>0.05, **p>0.01 and *** p>0.001 using multiple comparison with Ordinary One-Way ANOVA.

Furthermore, reduced RNA transcription post-irradiation was observed to different extent in various cell cycle phases in cells which were pre-depleted of HR factors CtIP, BRCA1 or RAD52, with the nascent transcription being more sensitive to BRCA1 depletion in S and G2 (Fig. 1f and Supp. Fig. 2a). These results are consistent with the recognized role of BRCA1 in transcriptional regulation of RNAPII, along with transcriptional activators such as p300/CBP(28). CtIP depletion negatively affected nascent RNA predominantly in the late S and G2, having less impact in the early S phase (Supp. Fig. 2a), where CtIP levels are still low. As expected, both BRCA1 and CtIP knockdown reduced ssDNA generation by resection in all cell cycle phases (Fig. 1g and Supp. Fig. 2b), while the CtIP-depleted cells showed a more pronounced resection defect in the G2 phase after DSB induction by irradiation, again coinciding with less nascent RNA (Supp. Fig. 2a, b).

Recently, it has been reported that RAD52 recruits BRCA1 in transcription-associated HR repair (8). Strikingly, the knockdown of RAD52 impaired the nascent RNA synthesis and DNA resection at IR-induced DSBs (Fig. 1e-g and Supp. Fig. 2a, b) but not significantly in non-damaged cells (Fig. 1e and Supp. Fig. 1b). Such nascent RNA and resection deficiencies associated with RAD52 depletion occur in all cell cycle phases upon irradiation (Supp. Fig. 2a, b), suggesting that both defects happen in a cell cycle-independent manner. This suggests that RAD52 plays a specific role in RNA transcription in response to DNA damage, at least during the first 60 minutes upon irradiation. Additionally, depletion of 53BP1, known for its role in promoting NHEJ, did not alter nascent RNA in either non- or irradiated-cells at either 30 or 60 minutes after irradiation (Supp. Fig.2 c, d). Taken together, our data show that nascent RNA transcription tightly coincides with DNA end resection in response to IR-generated DSBs, putting in context the relevance of transcription during different cell cycle phases and its alleged role in DNA repair choice upon irradiation.

### Nascent RNAs generated after DNA damage co-localize with DNA resection tracts

Next, we investigated whether the new RNA synthesis after DNA damage is required to facilitate DSB repair by HR. For this purpose, we developed a new technique to simultaneously observe nascent RNA and DNA resection tracts based on the principle of the nucleic acid (NA) fibers approach. Our technique, called R-SMART (RNA-SMART) to acknowledge a modification of the SMART assay (27), requires incubation with the RNA precursor 5-ethynyl uridine (EU) upon IR. To perform this analysis, Hela cells were treated with BrdU for 24 hours, followed by a 30-minutes EU pulse immediately after IR exposure, before stretching the NA fibers (Fig. 2a). The main objective of this technique is to quantify nascent RNA transcripts (through detection of the pulse-incorporated EU) only in DNA resection tracts. We measure the colocalization of EU-labeling and staining for BrdU under native conditions, allowing BrdU visualization in ssDNA, thus marking the DNA resection tracks (Fig. 2a, b). We observed a BrdU increased signal in fibers from cells exposed to 1 Gy and 5 Gy of ionizing radiation (Supp. Fig. 3a). Using the colocalization R-SMART technique, we detected an increased number of ssDNA fibers and their length in irradiated cells, using native conditions for anti-BrdU staining to visualize activation of DNA resection (Fig. 2b and Supp. Fig. 3b). Hence, the R-SMART technique can differentiate ssDNA among responses to different radiation doses (Supp. Fig. 3a) and can detect a deficiency in HR factors upon irradiation (Supp. Fig. 3b). Notably, profile analysis of BrdU-marked ssDNA resection tracts determined on the DNA fibers showed overlapping EU-labeled nascent RNA peaks, with colocalization of both accentuated in IR-treated cells (Fig. 2b-d). To better determine the impact of irradiation, we made a quantitative comparison of the overlap between nascent RNA and ssDNA in irradiated *versus* control cells. A more pronounced EU staining above the DNA resection tracts, seen in response to IR, suggested that nascent RNAs are linked with ssDNA generation (DNA end resection), the critical step of HR repair (Fig. 2c, d).

**Fig. 2.**
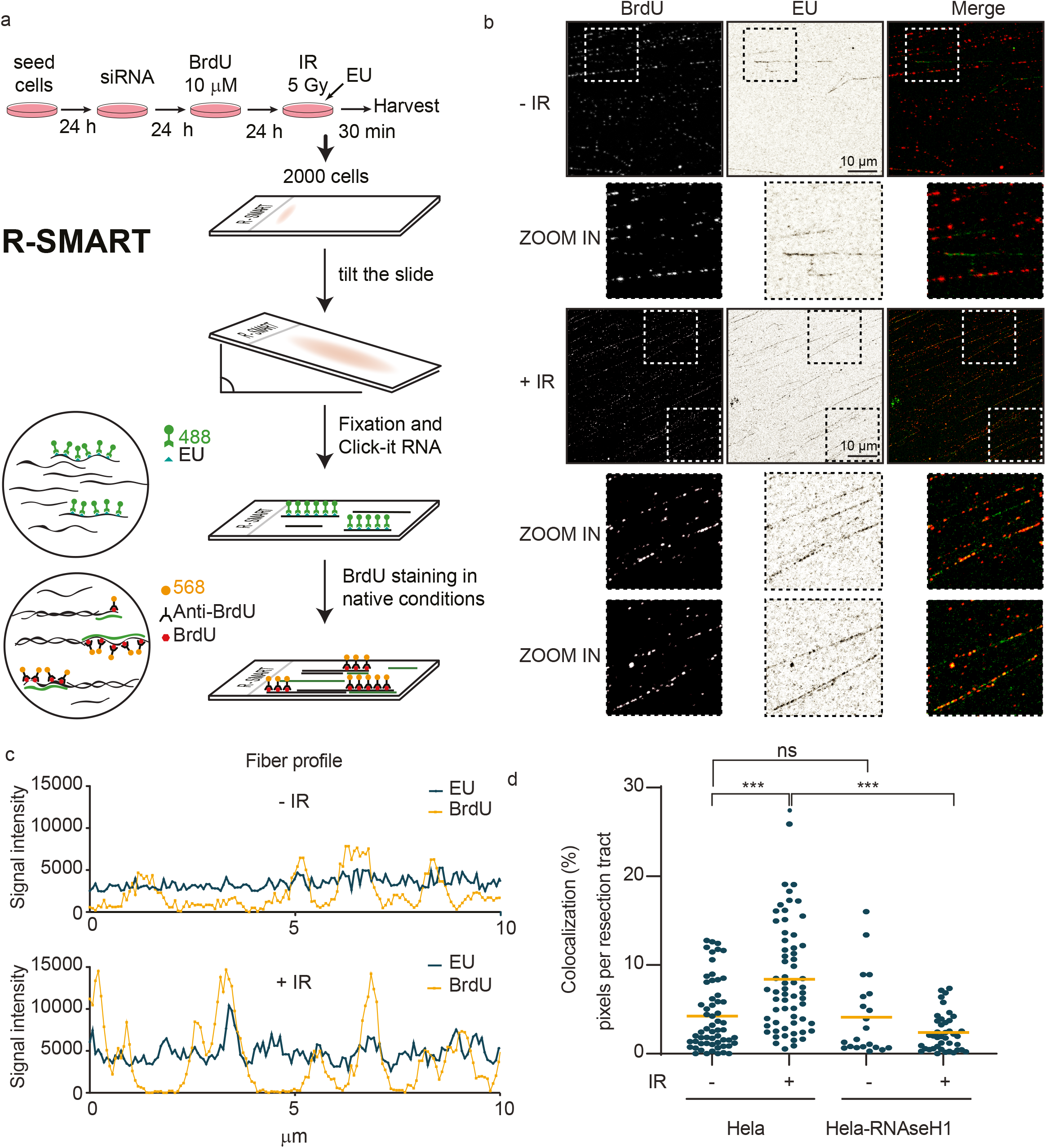
Nascent RNAs colocalize with DNA resection tracts in irradiated HeLa cells. **a** Schematic representation of the developed R-SMART technique. **b** Representative images of DNA resection tracts (black background) and nascent RNAs (white background) upon 5 Gy irradiation, using non-denaturing conditions for BrdU staining and EU labeling, respectively. Scale: 10um **c** Representative quantification of fibre profiles for EU and BrdU intensities from non-and irradiated Hela cells. **d** Dot graph shows percentage of BrdU and EU signal colocalization on resection tracts generated in non- and 5 Gy-irradiated Hela and Hela-RNAseH1 cells. At least 60 fields from 3 independent experiments were quantified. ns=non significant, *p>0.05, **p>0.01 and *** p>0.001 using multiple comparison with Ordinary One way ANOVA.

Based on these results, we would predict that the newly generated RNA at DSB regions would have high affinity and complementarity for the ssDNA stretches generated by resection during HR, thus likely creating DNA-RNA hybrid structures susceptible to degradation by RNAseH1 enzyme. This prediction was tested and confirmed, taking advantage of stable cell line HeLa-RNAseH1. HeLa cells were used in this section of our study due to the availability of a HeLa-derived cell line with overexpression of RNAseH1, a helpful model to assess any role(s) of RNA:DNA hybrids. RNaseH1 expression reduced the observed post-IR interaction between ssDNA and nascent RNA, confirming that ssDNA tracts are indeed likely covered or protected in some regions from degradation by nascent complementary RNA molecules (Fig. 2d). Altogether, these results support the notion that *de novo* RNA synthesis after DSB generation is closely linked to DNA resection during the early stage of DSB repair by HR.

### DNA damage stimulates RNA:DNA hybrid formation on DNA resection tracts

As RNAseH1 impacted the new RNA synthesis over ssDNA resection tracts in DNA-damaged cells, we next investigated whether such resection-associated RNA:DNA hybrids were detectable using the commonly employed S9.6 antibody (4). We took advantage of another optimized nucleic acid fibers-based technique, developed by us, called RL-SMART to visualize S9.6 staining in non-denatured nucleic acid fibers labelled and co-stained by antibody to BrdU (Fig. 3a and Supp. Fig. 4a). Using this technique, we detected a radiation-induced increase of RNA:DNA hybrids in nucleic acid fibers from cells exposed to 1 and 5 Gy doses (Supp. Fig. 4b), with up to 49,1% more RNA:DNA hybrids found in the irradiated samples (Fig. 3c). While only a fraction of RNA:DNA hybrids coincided with BrdU staining in the same track, the incidence of RNA:DNA staining colocalized with ssDNA tracts increased by 85% in irradiated cells (Fig. 3a, b). In parallel control cells, such RNA:DNA hybrid signal over ssDNA resection tracts was reduced when cells expressed ectopic RNAseH1 (Fig. 3a, b). RL-SMART profile analysis revealed that the RNA:DNA hybrid-staining peaks coincided with higher DNA resection staining peaks, indicating that the RNA-DNA hybrids indeed localize to the resection tracts generated upon irradiation (Fig. 3a, b). As expected, RNAseH1 overexpression in Hela cells abolished RNA:DNA hybrids on resection tracts (Fig. 3a, b). Quantification of RNA:DNA hybrids showed a 48% and 52% reduction of RNA:DNA hybrids seen in non-and IR-exposed Hela-RNAseH1 cells, respectively, compared with Hela control cells without ectopic RNAseH1 (Fig. 3c, d). Some residual RNA:DNA hybrids remained present despite RNAseH1 treatment, suggesting that some RNA:DNA hybrids are not entirely accessible and thus protected against degradation during DNA resection at DSBs and/or require other mechanisms to be resolved.

**Fig. 3.**
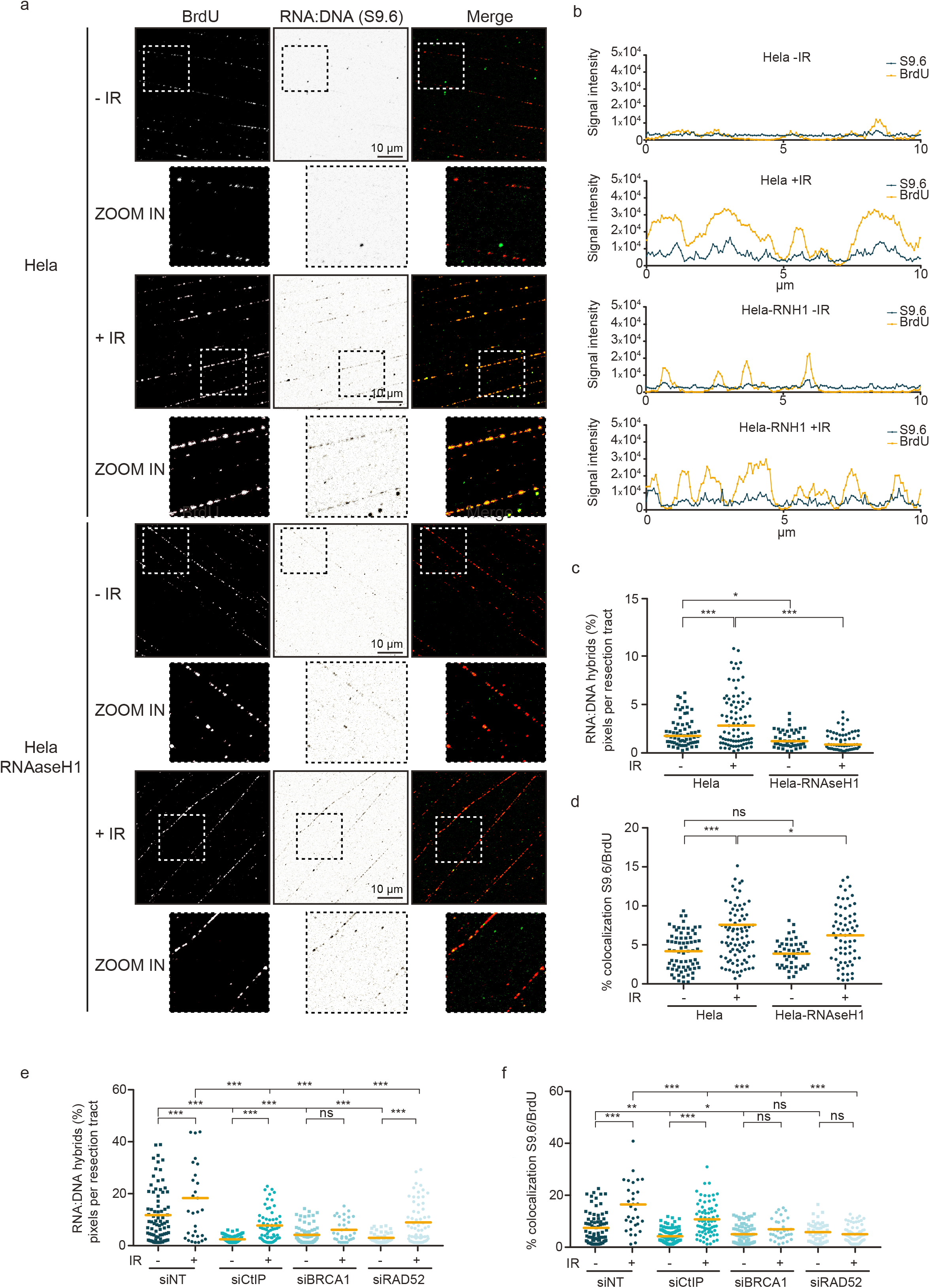
DNA damage generates high frequency of RNA:DNA hybrids on resection tracts. **a** Representative images showing ssDNA (resection tracts) and RNA:DNA hybrids upon non- and 5 Gy-irradiated Hela cells: non-denaturing BrdU staining and S9.6 staining, respectively. Scale 10μM **b** Representative quantifications of fiber profiles for Nascent RNA (EU) and RNA:DNA hybrids (S9.6) staining intensities from non- and irradiated Hela and Hela-RNAseH1 cells. **c** Dot graph shows percentages of RNA:DNA hybrids (S9.6) staining signal on resection tracts generated in non- and 5 Gy-irradiated Hela and Hela-RNAseH1 cells. **d** Dot graph shows percentages of RNA:DNA hybrids (S9.6) and ssDNA (BrdU) signal colocalization on resection tracts generated in non- and 5 Gy-irradiated Hela and Hela- RNAseH1 cells. **e** Hela cells transfected with siRNA against CtIP, BRCA1, RAD52 and NT (Non-Target) for 48 hours, were assessed for RNA:DNA hybrids (S9.6) quantification by RL-SMART as in (c). **f** Hela cells transfected with siRNA against CtIP, BRCA1, RAD52 and NT (Non-Target) for 48 hours were assessed for RNA:DNA hybrids (S9.6) and ssDNA (BrdU) and quantify the percentage of colocalization as (d). **b-f**, at least 30 fields from each of ≥ 3 independent experiments were quantified. ns=non-significant, *p>0.05, **p>0.01 and *** p>0.001 using multiple comparison with Ordinary One Way ANOVA.

Next, we studied the role of HR factors in RNA:DNA hybrid formation during DNA resection using the RL-SMART technique. Consistently with the above results, when Hela cells were depleted for CtIP, BRCA1 and RAD52 proteins, respectively, we observed more than 50% decreased RNA-DNA hybrid formation in all cases (Fig. 3e-f). These results most likely reflect the decreased formation of nascent RNA during the DNA repair by HR (Fig. 3e-f). The extent of RNA:DNA hybrids were reduced in irradiated and non-irradiated cells, the latter likely reflecting repair of endogenous DNA damage, possibly caused by cancer-associated endogenous replication stress and the ensuing DSBs (26), the repair of which is also affected by depletion of these DDR factors. Interestingly, irradiated CtIP-depleted cells still generated detectable RNA:DNA hybrids, albeit reduced at 61,9% in comparison with the irradiated CtIP-proficient controls (Fig. 3e-f). Cells depleted of RAD52 also showed a reduction in RNA:DNA structures, the incidence of which can increase after irradiation (Fig. 3e), although in this scenario, colocalization with DNA resection tracts did not increase (Fig. 3f). BRCA1 protein was also required for RNA:DNA hybrid formation after DNA damage, in terms of both the overall extent and colocalization with DNA resection tracts (Fig. 3e-f). Taken together, these results indicate that new transcript RNAs generate RNA:DNA hybrids using 3^’^-5’ssDNA, thereby providing an additional level of regulation over DNA resection, suggesting a potential new role of CtIP and BRCA1 in DNA repair pathway choice.

### RNAPII inhibition impairs the recruitment of HR proteins

Given our results so far and the fact that RNA polymerase II (RNAPII) gets recruited to DSBs (21,23), we hypothesized that RNAPII accumulation and newly produced RNA could be involved in choosing between the two main DSB repair pathways. To investigate this hypothesis, we used a THZ1 compound, a potent inhibitor of CDK7, indeed of RNAPII-mediated RNA transcription (29), to assess the impact of RNA synthesis inhibition on recruitment of DDR proteins to DSB. We monitored phosphorylation of Serine 5 on CTD repeats of RNAPII by CDK7, a modification which is essential for transcription initiation, and demonstrated specific inhibition of phosphorylation of this residue upon THZ1 treatment (Supp. Fig. 5a). Next, we analyzed the recruitment of the main DSB repair factors tagged with the green fluorescence protein (GFP) to microlaser irradiation-created DNA damage tracts. Given that CtIP is a key protein for HR-mediated DSB repair, playing a major role in the activation of DNA resection (30-32), and our present data showing the impact of CtIP on radiation-induced RNA:DNA hybrid formation, we evaluated CtIP-GFP dynamics on DNA damage sites in THZ1-treated cells (Fig. 4a-b). RNAPII inhibition, using THZ1 treatment, reduced the CtIP-GFP recruitment to DSBs during the initial 10 minutes after microirradiation (see also the supplementary video 1). We tested CtIP recruitment in cells treated with DRB, a known RNAPII inhibitor, observing similar CtIP recruitment pattern that THZ1 treatment (Supp. Fig. 5b). These results suggested that functional RNAPII is required for the initiation of CtIP-dependent DNA resection. It is well known that HR deficiency leads to DSB repair shift towards NHEJ repair (13,14). Notably, studying 53BP1-GFP kinetics in the presence of RNAPII inhibitor, we found that 53BP1 recruitment was more abundant and faster than in control vehicle-treated cells (Fig. 4c-d), suggesting that DNA resection deficiency due to RNAPII inhibition favours the NHEJ pathway. DSB repair shift towards NHEJ is created around 4 minutes (240 sec) upon irradiation, whereas the impairment of CtIP recruitment occurs already during the first 2 minutes after laser microirradiation. When other GFP-tagged DDR proteins were evaluated, we found that MRE11, another key resection protein, also displayed an acutely impaired recruitment to DSBs under THZ1 treatment (Fig. 4e). These data suggest that CtIP and MRE11, as key DNA resection factors, require RNAPII activation at DSB sites to promote the DNA resection machinery assembly and thereby promotion of the HR repair pathway. BRCA1 and RPA recruitment were also reduced upon RNAPII inhibition but slightly later than CtIP and MRE11, suggesting that these factors help DNA resection processing at a later step rather than at the initiation. Indeed, BRCA1 is essential to regulate resection speed and RPA protects ssDNA after resection (25) (Fig. 4f, h). Furthermore, recruitment of RAD52, a protein shown to promote transcription-coupled HR repair, to the DNA repair complex was also impaired under THZ1 treatment (Fig. 4g), consistently with the reported impact of the RNAPII inhibitor DRB (8). Taken together, our data show that RNAPII activity supports efficient recruitment of HR repair factors, and inhibition of RNAPII favours NHEJ repair through preferential recruitment of 53BP1 early in the chromatin response to DSBs.

**Fig. 4.**
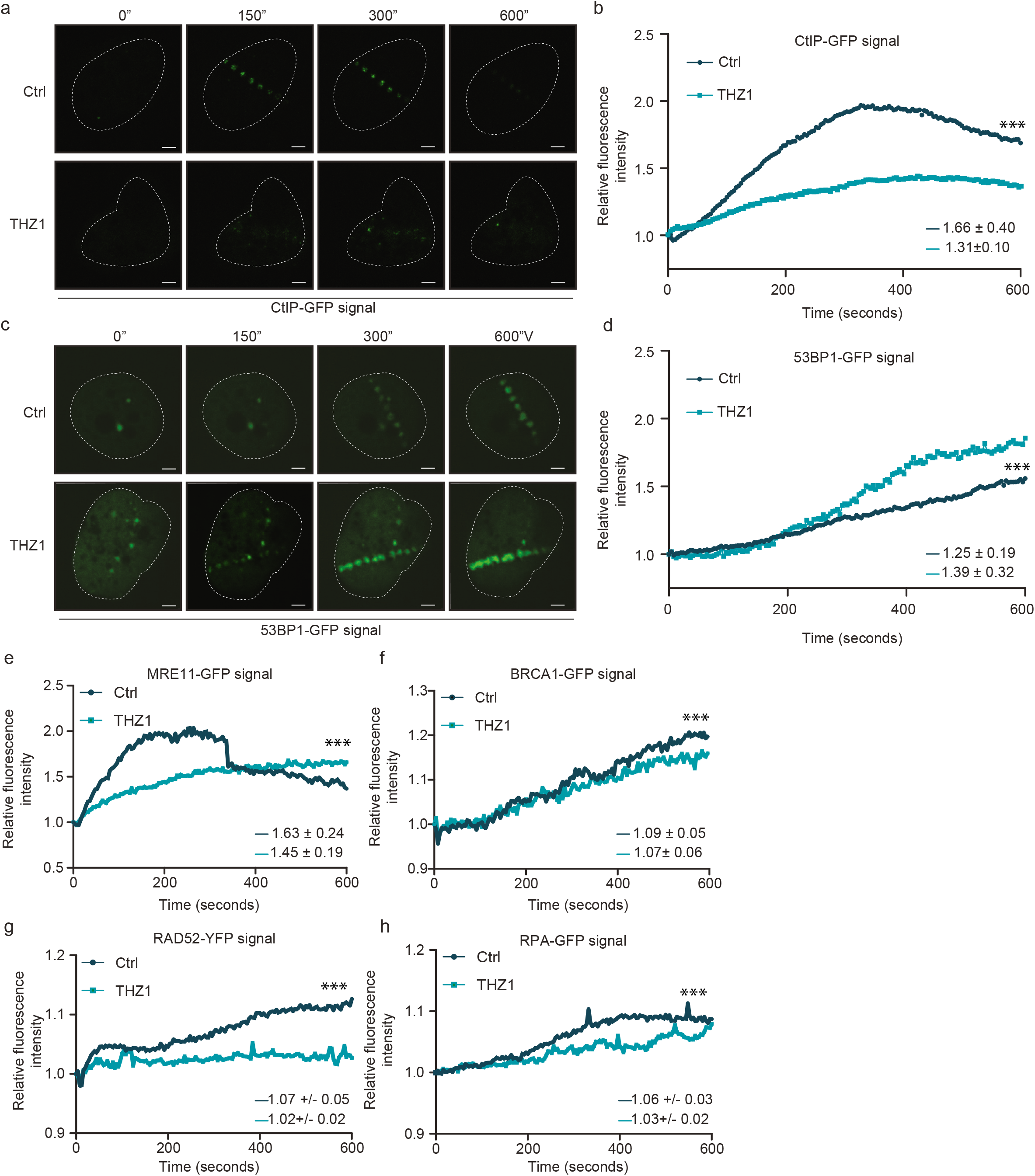
Inhibition of RNAPII impairs DDR protein recruitment to DSBs. **a** Representative time-lapse images of CtIP-GFP recruitment to DNA damage in RNAPII - inhibited (THZ1, 1 µM during 2 hours) and DMSO as control during 600 seconds post microlaser irradiation in U2OS cells. Scale bar: 1 μm **b** Effect of THZ1 treatment (1 µM during 2 hours) on the kinetics of CtIP-GFP recruitment to DNA damage by measuring relative fluorescence intensity of CtIP-GFP during 600 seconds post microlaser irradiation in U2OS cells. The analysis represents the average on n=30 (Ctrl) and n=15 (THZ1) nuclei from 3 biologically independent experiments. **c** Representative images of 53BP1-GFP recruitment to DNA damage after THZ1 treatment (1 µM during 2 hours), assessed as in (**a)**. Scale bar: 10 μm **d** Kinetics of 53BP1-GFP intensity in U2OS cells treated with DMSO or THZ1 (1 µM during 2 hours) prior microlaser irradiation, measured as in (**b)**. The analysis represents the average on n=17 (Ctrl) and n=18 (THZ1) nuclei from 3 biologically independent experiments. **e-h** Kinetics of recruitment to laser-induced DNA damage, for DDR factors MRE11 (**e**), BRCA1 (**f**), RAD52 (**g**) and RPA (**h**) treated or not with RNAPII inhibitor (THZ1, 1 µM during 2 hours) prior microlaser irradiation, and measured as in (**b)**. The analysis represents the average of nuclei from 3 biologically independent experiments. MRE11, n=31 (Ctrl) and n=29 (THZ1); BRCA1, n=18 (Ctrl) and n=14 (THZ1), RAD52, n=26 (Ctrl) and n=8 (THZ1), RPA, n=17 (Ctrl) and n=16 (THZ1). b-d,e-h, two-tailed paired t-test was analyzed for all kinetics. *** p>0.001. In the graphs the mean of mobile fractions and the ±SD are shown for each sample.

### Functional RNAPII shifts the DSB repair balance towards HR

Next, we wished to extend further the concept emerging from our present results, namely that new RNAPII-mediated transcription at DSB sites is required to direct the cell’s choice of DSB repair process by promoting accrual of essential HR factors. First, we demonstrated that recruitment of RPA, which is required to protect ssDNA stretches created during DNA end resection, diminished at DSBs during the first 10 minutes after microirradiation (Fig. 4h). Next, we wanted to know whether cells exposed to ionizing radiation and treated with THZ1 alter their choice of DNA repair mode in the longer term. We exploited RPA foci formation as a DNA resection marker until two hourspost irradiation, and found that such cells became indeed defective in their ability to form foci of phosphorylated RPA (Supp. Fig. 6a). However, another RNAPII inhibitors such as DRB and α-amanitin did not impair phosphorylated RPA foci formation (Supp. Fig. 6a). When testing the kinetics of other DDR factors to generate IR-induced foci in THZ1-treated cells, we observed enhanced accumulation of 53BP1 to DSB-flanking chromatin, associated with an increase of nuclear foci formation of 53BP1 until 2 hours after irradiation under THZ1 treatment (Fig. 5a). 53BP1 foci accumulation in THZ1-treated cells was almost 5-fold increased during 1 hour even without any exogenous genotoxic insult, suggesting that under RNAPII inhibition conditions, endogenously arising DNA lesions are preferentially repaired by the error-prone NHEJ pathway.

**Fig. 5.**
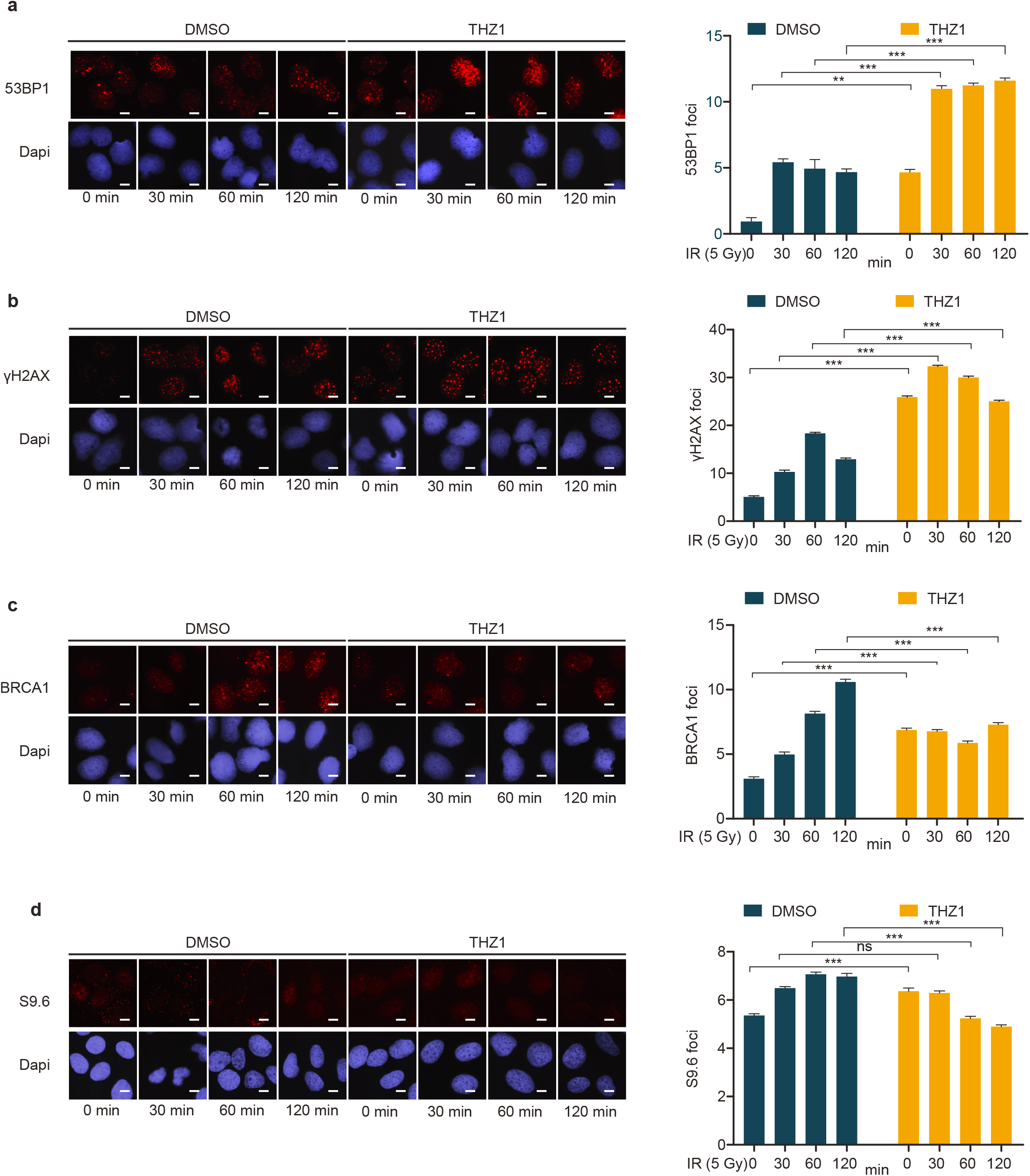
Defective recruitment of HR factors after RNAPII inhibition favors NHEJ repair. **a** Representative images of 53BP1 foci upon 5 Gy-irradiation in U2OS cells treated with DMSO or THZ1 (1 µM during 2 hours), (left). Bar graph for 53BP1 foci quantification at 0, 30, 60 and 120 minutes after irradiation (right). **b** Same as in (**a**), but measuring γH2AX foci **c** Same as in (**a**), but measuring BRCA1 foci **d** Same as in (**a**), but measuring RNA:DNA hybrids foci using S9.6 antibody **a-d**: At least 500 cells from ≥ 3 independent experiments were quantified. ns=non-significant *p>0.05, **p>0.01 and *** p>0.001 using multiple comparison with Ordinary One-Way ANOVA. Scale bar: 10μm

Furthermore, we tested RIF1, as a 53BP1 interactor when phosphorylated by ATM to promote NHEJ, in THZ1-treated cells and showed that RNAPII inhibition increased this marker of NHEJ upon IR as well (Fig. Supp 6b). Also, we tested whether other RNAPII inhibitors such as DRB or a-amanitin have the same effects as the specific inhibitor THZ1. Surprisingly, THZ1 showed a higher ability to impair HR and promote NHEJ than the other broader-specificity inhibitors tested, when monitoring foci formation by different DSB response factors (Fig. 5a-c and Supp. Fig. 6).

The THZ1-inhibited cells showed more unresolved DNA damage associated with enhanced DNA damage marker formation before and after irradiation (Fig. 5a-c), demonstrating that RNAPII inhibition play an active role in endogenous DNA damage. Furthermore, foci formation by BRCA1 and RAD51, factors involved in early and late HR steps, respectively, was impaired at multiple time points post-irradiation in the THZ1-treated cells (Fig. 5c and Supp. Fig. 6c). Consistent with our previous data using the RL-SMART assay and the contribution of RNA:DNA hybrids to proper DNA end resection emerging from our present results, we observed that RNA:DNA hybrid foci were significantly reduced after treatment with THZ1 until 2 hours after irradiation (Fig. 5d). Thus, we conclude that nascent RNA synthesis is an essential step to promote DSB repair *via* HR and that formation of RNA-DNA hybrids at DSBs provides an intermediate structure required to carry out the early phase of HR.

### CtIP is required to re-initiate transiently paused RNA transcription after DNA damage

Apart from indicating that the observed DSB-associated transcription shifts the DSB repair pathway choice towards HR, our results also suggested that the DNA resection factors could be promoting transcription to facilitate 5^’^-3’ single-strand DNA degradation. Indeed, while CtIP and BRCA1 are critical for DNA resection during HR repair, these factors are also known to function as transcription factors (33,34), albeit the molecular basis for the latter role and how it may be coordinated with the DNA resection role during DSB repair, are not fully understood. Thus, we hypothesized that CtIP could regulate transcription at DSBs, thereby allowing other DNA resection proteins to be recruited. To address this possibility, we first tested *de novo* RNA transcription in CtIP-deficient cells before and at early time points after DNA damage. Run On technique is a complementary approach to EU-labelling, we allow us to measure RNA transcription at 7,15 and 30 minutes upon irradiation. Depletion of CtIP did not significantly alter RNA transcription in non-damaged U2OS cells when assessed by BrUTP incorporation into nucleoplasmic nascent RNA (Fig. 6a-b). As expected, global transcription decreased upon DNA damage in a CtIP-independent manner. However, BRCA1-deficient cells showed reduced RNA transcription under non-irradiated conditions (Supp. Fig. 7a, b), suggesting a possible replication- and/or transcription-associated function of BRCA1 independent of exogenous DSBs.

**Fig. 6.**
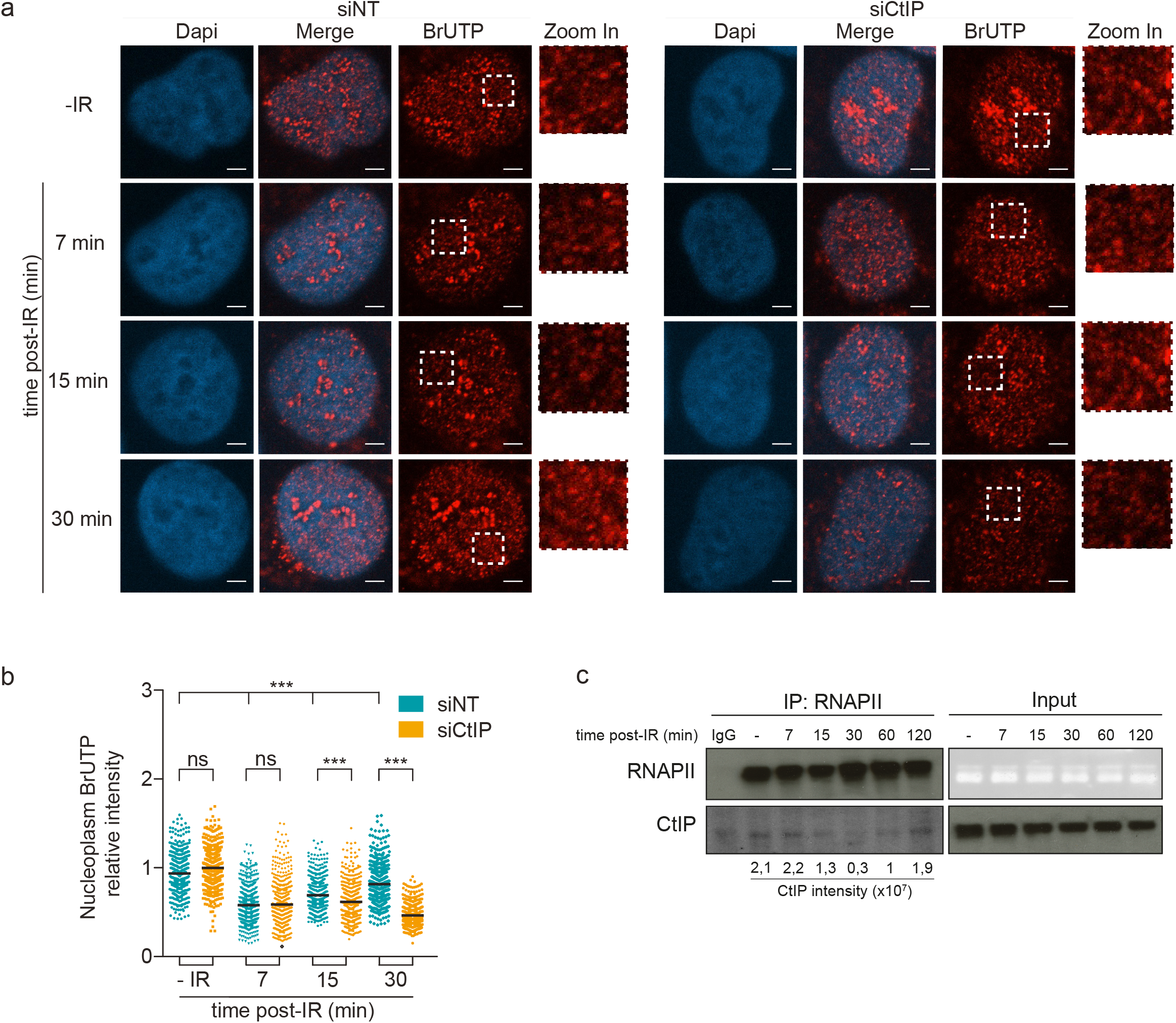
DNA resection deficiency prevents transcription re-start after DNA damage. **a** Representative images of *de novo* transcription: BrUTP incorporation into nascent RNA at 7, 15 and 30 min post-IR (5 Gy) in CtIP-depleted U2OS cells. Zoom in pictures are shown. Scale bar: 1μm. **b** Dot graph shows nucleoplasm quantification of BrUTP incorporation under the experimental conditions cued in (**a)**. At least 200 cells from each of ≥ 3 independent experiments were quantified. ns=non-significance *p>0.05, **p>0.01 and *** p>0.001 using multiple comparison with Ordinary One-Way ANOVA. **c** RNAPII immunoprecipitation from extracts of non- and irradiated U2OS cells at 7, 15, 30, 60 and 120 min post-IR (5 Gy). Immunoblot detection of CtIP and RNAPII proteins in immunoprecipitated and input samples, as indicated. Quantification of CtIP co-immunoprecipitated with RNAPII is shown below.

Interestingly, our time-course experiments suggested that CtIP and also BRCA1 are required for the timely recovery of global cellular transcription after 30 minutes post-irradiation, while being indispensable for the generation of nascent RNA associated with DSB repair very early after irradiation (Fig. 6a-b and Supp. Fig. 7a-b). In this context, we also assessed CtIP interaction with RNAPII upon DNA damage to examine whether CtIP might regulate the transcription associated with DSB repair. Interestingly, CtIP interacted with RNAPII in non-irradiated as well as irradiated cells, yet such interaction was reduced during the initial 15 and 30 minutes post-IR, while recovering to pre-irradiation levels at 60 and 120 minutes after irradiation (Fig. 6c). Taken together with the other results of this study, we suggest that the CtIP plays a direct, dual role in both the initiation phase and later progression of transcription-coupled DNA resection in DSB repair by the HR pathway.

## Discussion

One of the major recent advances in biomedicine has been the realization that diverse forms of RNA are intimately involved in a much broader range of fundamental biological processes than traditionally thought. RNAs and RNA:DNA hybrid structures, commonly referred to as RNA:DNA hybrids, have been widely implicated in physiological mechanisms and molecular pathogenesis of grave diseases such as cancer. Among the emerging new roles of RNA is the causal involvement in various aspects of genome integrity maintenance, including repair of DNA DSBs, arguably the most hazardous genotoxic lesions (8,23,35). Indeed, assembly of functional RNAPII and non-coding RNA transcripts at DSBs can promote the chromatin DDR foci formation, at least in part by driving liquid-liquid phase-separation formation condensates of 53BP1, a factor involved in the regulation of the more error-prone DSB repair by NHEJ (23). On the other hand, the outstanding issues of whether and mechanistically how are DSB-associated RNAs involved in the choice between NHEJ and the error-free HR pathways have remained unsolved, despite recent studies on transcription-coupled DNA repair raised new questions about the specific role of RNA in DSB repair (11,12,36,37). Complementary to our data presented here, and two years after the submission of our present manuscript, it has been published that RNAPIII can create nascent RNA at DSBs, promoting assembly of the HR machinery (24). However, the latter work dismissed similar involvement of RNAPII, based on the observation that one of the RNAPII subunits, RPB1, was not detected as recruited on the laser-generated DSB stripe, and consequently exclusively focusing only on the RNAPIII subunits (24).

In this context, our present study complements and advances these efforts by providing a conceptual framework for, and mechanistic insights into, the role of nascent RNA transcripts and RNA:DNA hybrids at DSBs, as critical factors promoting DNA end resection and hence the upstream steps for the decision of human cells to repair DSBs by HR (see Fig. 7 for our proposed model). Indeed, our data demonstrate that RNAPII plays an essential role in this process. A vital prerequisite for addressing this issue was the development of our two new single-molecule nucleic acid analysis methods that we present here: R-SMART and RL-SMART. These methods allowed us to assess the RNA-related DSB-repair events upon irradiation of human cells with a clinically relevant dose of 5Gy, in an unbiased, pan-genomic manner. Indeed, ionizing radiation generates global DNA damage in both transcriptionally active and silent genomic regions. Based on the current knowledge, we suggest that the bulk of such random IR-induced DSBs occur in otherwise non-transcribed chromatin because over 90% of the human genome is free from either protein-coding genes (16) or non-protein-coding yet transcribed elements (38). It is known that the global RNA transcription activity becomes inhibited upon DNA damage, and factors including ATM and cohesion contribute to this mechanism (17,39). Also, a recent study demonstrated nascent RNA break-induced transcriptional suppression using a single-molecule transcription reporter upon controlled induction of DSB (9). In our present study, RNA transcription silencing occurred within the initial 30 minutes upon irradiation, consistent with the published reports (9, 17, 48). However, we show that nascent RNA formation is recovered by 60 minutes, and it is already increased by 30 min post irradiation, compared to non-irradiated cells (Fig. 1 and Fig. 5), probably as a consequence of nascent RNA generated at DSB sites combined with the regular activity of RNAPII in the transcriptionally active genomic regions, the latter responsible for the nascent RNA detected also in the mock-treated cells. Within the initial 30 minutes post irradiation the choice of DSB repair pathway is made, a process in which our results show the active regulation of RNAPII-mediated transcription at DSB sites is essential. At the same time, regulation of DSB-associated nascent RNA could depend on the genomic context (9,35) as well as on the nature of the genotoxic insult and the type of the ensuing DNA lesions. Notably, our results also challenge the notion that HR-mediated repair might be limited to only those DSBs present in genomic loci that are transcriptionally active under physiological, non-stressed conditions (8). The results from the R-SMART technique allowed us to conclude that nascent RNAs actively generated during DSB repair are linked to ssDNA resection tracts, implicating new RNA synthesis during HR (Fig. 2).

**Fig. 7.**
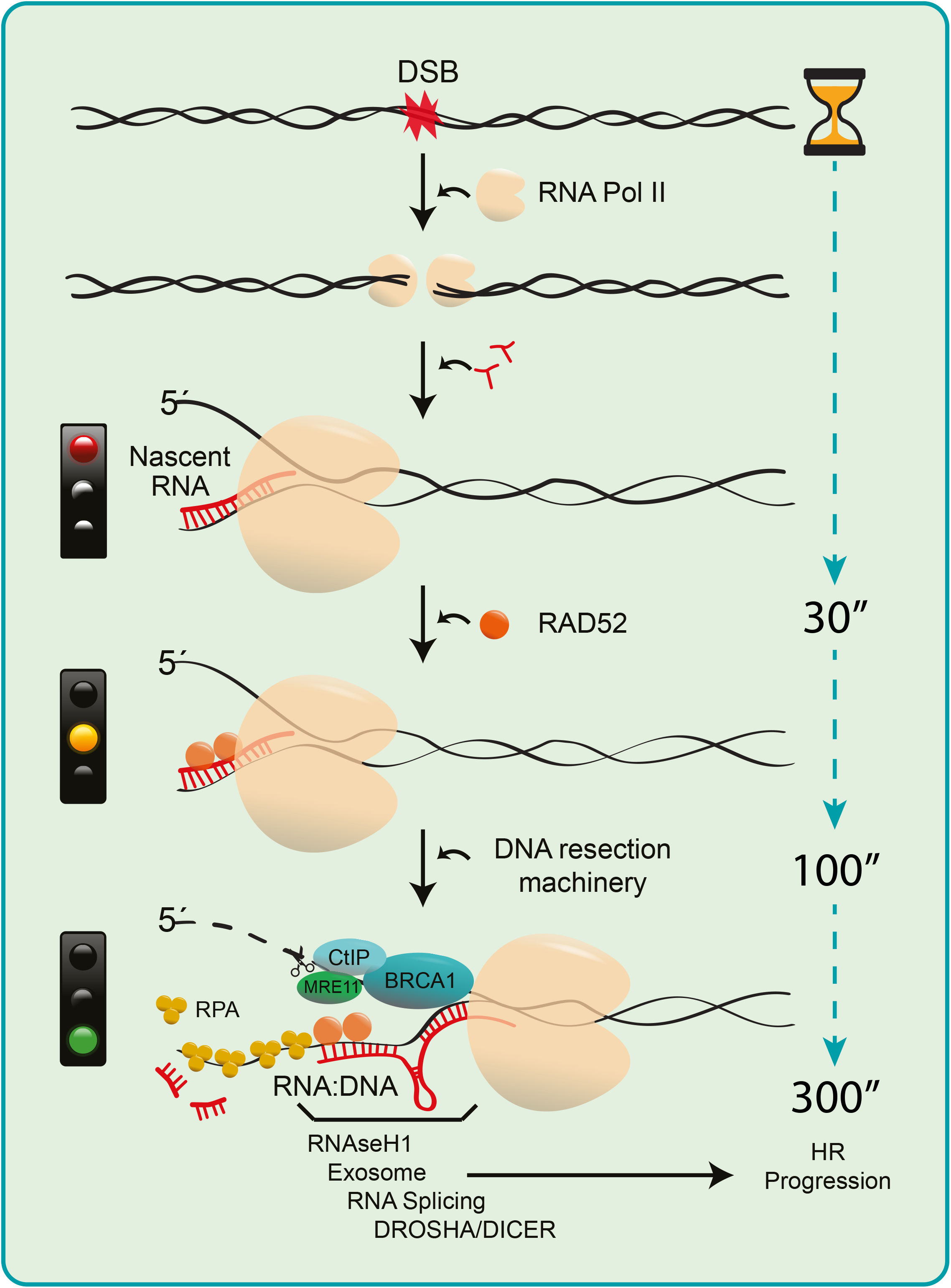
Schematic model of the regulation and role of RNAPII-generated nascent RNA to guide DNA end resection and DSB repair by HR. Transcriptionally active RNAPII is more prominent in S-G2 cell cycle phases (showed in Fig.1) when HR is known to repair DSBs. RNAPII recruitment to DSBs, that involves the pre-initiation complex (PIC) (23), favours nascent RNA transcription, leading to generation of small DNA:RNA hybrid structures that cause RNAPII pausing (showed in Fig.2 and 3). RAD52 and XPG are capable of rescue the transiently paused RNAPII activity (8), allowing for 5’DNA strand displacement. Experimental inhibition of transcription impairs recruitment of the DNA resection factors CtIP, MRE11 and BRCA1 (showed in Fig.4 and 5) to DSBs. MRE11 initiates 5’strand degradation, while the CtIP-BRCA1 axis is essential to regulate the speed of resection (23) by controlling transcription upon DNA damage (showed in Fig.6). We show that CtIP and BRCA1 are key factors promoting and/or re-starting the locally paused RNAPII-mediated transcription. These upstream events guide the DSB repair choice towards HR through initiation and progression of DNA end resection, in a feedback loop in which proper CtIP and BRCA1 recruitment are stimulated by the RNAPII at the DSB site, at least in part through complex formation between CtIP and RNAPII. Additional proteins and auxiliary processes such as exosomes (58) RNA splicing (50) and Drosha/Dicer (20) contribute to regulation of DNA:RNA hybrids along the initiated HR pathway to adjust this process in a cell context- and time-dependent manner to safeguard genomic integrity.

Furthermore, using the related RL-SMART approach, we observed that *de novo* transcription around DSB sites promotes the formation of RNA:DNA structures sensitive to resolution by RNAseH1 (Fig. 3) and possibly by other proteins such as Senataxin (19) or XPG (8). While the accumulation of some RNA:DNA hybrids can undermine genomic stability, RNA-DNA hybrids are also required for efficient DSB repair (4,7). RNA:DNA hybrids commonly form when the RNAPII pauses during transcription (4), thereby, in the context studied here, allowing recruitment of DDR factors to the DNA break. At the same time, RNA:DNA hybrid generation during the DSB-flanking RNA synthesis is primarily affected by depletion of CtIP or BRCA1 during the first 30 minutes after DNA damage. (Fig. 3). These results suggest that RNAPII recruitment to DSBs and acute bidirectional transcription (9) results in the first round of RNA transcripts forming RNA:DNA hybrids and DNA resection factors becoming recruited after RAD52-dependent resolution of these hybrid structures (Fig. 7). Paused RNAPII activity needs to be re-launched after incorporation of DNA resection machinery into the DNA repair complex, and we show that efficient recruitment to DSBs of CtIP, a factor involved in DNA resection, requires active RNAPII (Fig. 4) as well as MRE11 for the early step of transcription-associated DSB repair. Furthermore, recruitment of HR factors such as RPA, BRCA1 and RAD52 to DSBs was also impaired after RNAPII inhibition by THZ1. Under such RNAPII-inhibition conditions, recruitment of the NHEJ-related 53BP1 protein was initially favoured on the DSB-flanking chromatin, suggesting that proficient nascent RNA transcription shifts the balance of DSB pathway choice towards HR. In contrast, RNAPII inhibition early after IR favours the recruitment of 53BP1 and RIF1, factors implicated in NHEJ. Indeed, PBAF-mediated transcription silencing flanking DNA breaks contributes to NHEJ (40), and H4 deacetylation, which is associated with repression of transcription, favours 53BP1 foci formation early upon DNA damage (41).

We also show that RNA synthesis inhibition impairs both the formation of RNA:DNA hybrids in the proximity of DSBs and HR repair, consistently with recent studies in yeast (7) and studies of mammalian cells that however focused only on transcriptionally active regions of the genome (8,19,20). Interestingly, while CtIP is essential to initiate DNA resection, CtIP’s precise mechanistic role in DNA resection has remained unclear. We demonstrate a so-far unsuspected function of CtIP in the re-starting of RNA transcription upon DNA damage, as a step necessary to carry out DNA end resection properly, and reflecting a direct CtIP-RNAPII interaction reported here, as a prerequisite for this role. Related to this, CtIP’s partner in DNA resection, BRCA1, was also required for the transcription associated with DSB repair. Our data suggest that CtIP, BRCA1 and their interplay are essential for re-launching the transiently paused activity of RNAPII and further progression of DNA resection (Fig. 7).

Overall, such extended transcription-promoting role of CtIP and BRCA1 is broadly consistent with, and complements, previous reports (34,42,43). Furthermore, the fact that CtIP prevents excessive RNA:DNA hybrid accumulation after DNA damage (44), is consistent with the notion we propose here, namely that DNA resection is delicately regulated by *de novo* RNA transcription that relies on the BRCA1-CtIP axis. BRCA1 deficiency increases global R-loop formation (45) related to its multiple roles in diverse stages of the cell cycle, under replication stress and in DNA repair. In our study, we focused on the BRCA1-CtIP axis function on DNA end resection. In the absence of these HR factors, the abundance of RNA:DNA hybrids locally decreases on the DSB resection tracts (Fig. 3). The latter result could be linked with the role of BRCA1 in the re-activation of transcription elongation upon DNA damage (46) or the requirement of BRCA1/SETX complexes to mitigate R-loop-driven DNA damage (47).

Taken together with the current knowledge, our data, therefore, suggest the model (Fig. 7) in which RNAPII is actively recruited to the DSBs (21,23), nascent RNA transcription by RNAPII induces the local formation of RNA:DNA hybrids that trigger RAD52 incorporation to resolve these structures (8), together with RNAseH1 recruitment upon RPA coating of ssDNA(48). RNAPII pausing occurs as a regulatory step in transcription, allowing time and local environment to resolve the DNA-RNA and provide a displaced 5^’^-3’ssDNA available to recruit the resection machinery. CtIP, and its interaction with BRCA1(27,34), promotes RNA transcription, permitting 5’-3’degradation by exonucleases proteins such as MRE11(49). Regulation of RNA transcription through the CtIP-BRCA1 axis, removal of RNA:DNA hybrids, secondary RNA structures, RNA degradation and lncRNA for splicing machinery (50), exosome (51), and Dicer-Drosha (21) complexes may all contribute to managing the later steps of HR, allowing the accrual of the RAD51-BRCA2 (52) cascade whose presence and the ensuing resolution of these various structures could be essential to avoid genome instability.

In conclusion, nascent DSB-flanking RNAs and their local dynamics are essential for proper guidance of the cell’s decision to repair the potentially lethal DSB lesions *via* the faithful HR pathway, thereby safeguarding homeostasis and minimizing the development of pathologies, including cancer. Moreover, as cancer cells feature an enhanced load of endogenous DSBs (53-55), our present concept has also implications for the emerging cancer treatments by transcription inhibitors (56,57). This strategy likely exploits tumour cells’ vulnerability, whose survival and proliferation are ‘addicted’ to altered checkpoint signalling and dependent on repair mechanisms dealing with the excessive burden of chromosomal breaks.

## Supporting information

CtIP recruitment to DNA damage

## Acknowledgements

Thanks to Ekaterina Petrovicheva for technical assistance.

## Author contributions

D.G-C primarily conceived and designed the study in discussions with G.P, D.A-M and J.B. D.G-C performed most of the experiments. G.P carried out co-immunoprecipitation and run-on assays. D.A-M helped with figures and experimental design. C.D evaluated assay outcomes using ScanR analysis. D.G-C and J.B. wrote the manuscript, and J.B. supervised the study. All authors discussed and interpreted the data, and approved the manuscript.

---

**Table 1:** Antibodies used:

| **ANTIBODIES AGAINST** | **COMPANY** | **CATALOGUE** | **HOST** |
| --- | --- | --- | --- |
| **BRDU** | APBiotech | RPN20AB(RPN202) | Rat |
| **BRDU** | BD Biosciences | 347580 | Mouse |
| **S9.6** | keraFAST | ENH001 | Mouse |
| **53BP1** | millipore | MAB3802 | Mouse |
| **H2AX-S139** | Millipore | 16-193 | Mouse |
| **H2AX-S139** | Cell Signaling | S2577s | Rabbit |
| **BRCA1** | Santa Cruz | sc-6954 | Mouse |
| **MOUSE ANTI-RNAPOL II** | Abcam | ab817 | Mouse |
| **S5P RNAPOL II** | Abcam | ab5131 | Rabbit |
| **CTIP** | active motif | 61141 | Mouse |
| **RAD52** | Santa Cruz | sc-365341 | Mouse |
| **ALPHA-TUBULIN** | Gene Tex | GTX628802 | Mouse |
| **S4/S8 RPA** | Bethyl | A300-245A | Rabbit |
| **RAD51** | Abcam | ab63801 | Mouse |
| **PCNA** | immuno concepts | 2037 | Human |
| **RIF1** | Bethyl | A300-569A | Rabbit |

## Supplementary Figure Legends

**Suppl. Fig. 1.**
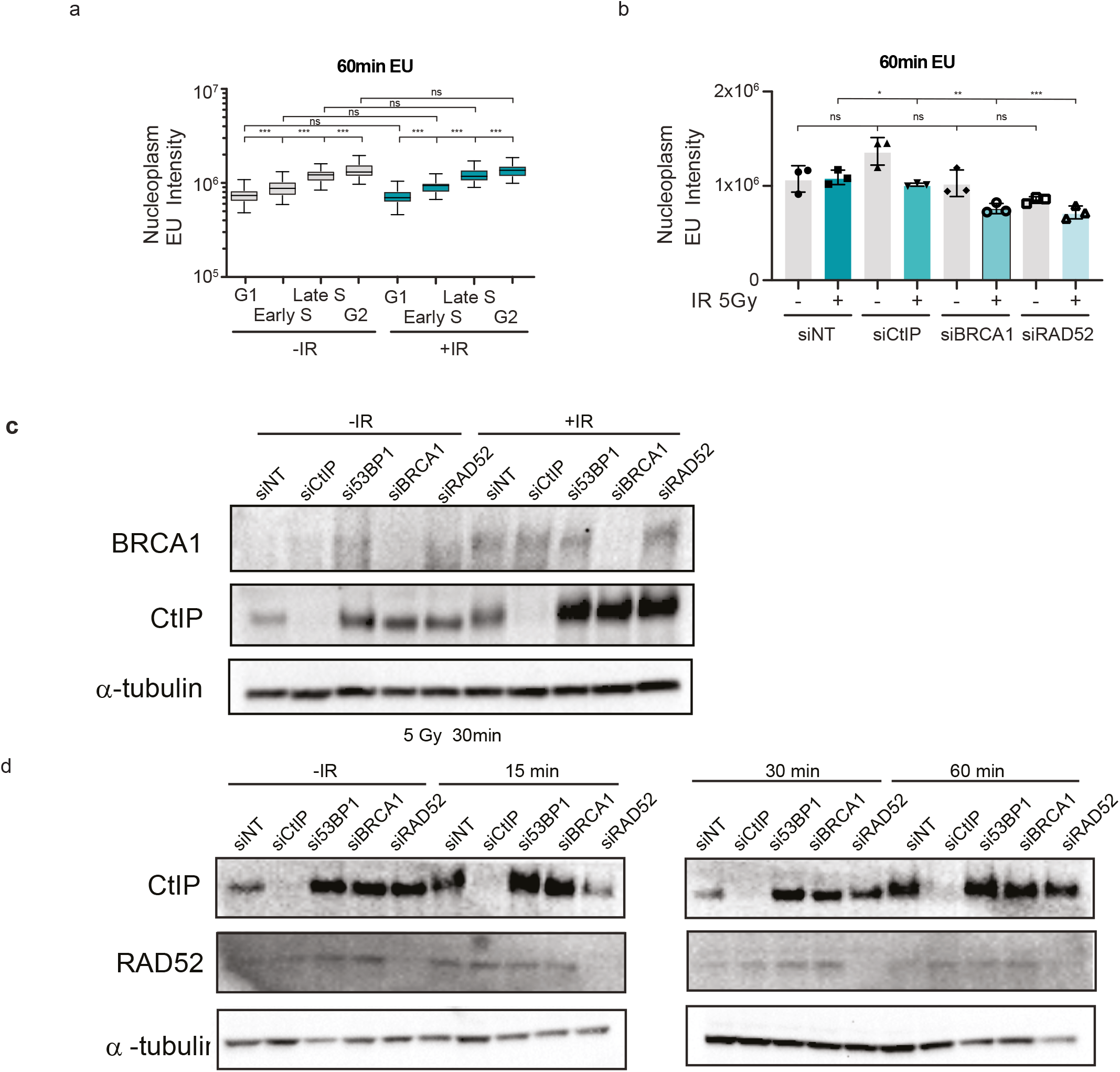
**a** Graph shows nucleoplasm EU intensity in different cell cycle phases in non- and irradiated-U2OS cells. EU component was added upon DNA damage using 5Gy in irradiated cells and labelled for 60 minutes in both cellular conditions. Mean value of 500 cells are represented and 95-5 percentile error bars. At least, 2 independent experiments were quantified. ns=non-significance *p>0.05, **p>0.01 and *** p>0.001 using multiple comparison with ordinary One-Way ANOVA. **b** Histogram represents nucleoplasm EU intensity in siRNAs against indicated genes in non-and irradiated-cells. EU component was added upon DNA damage using 5Gy in irradiated cells and labelled for 30 minutes in both cellular conditions. Showing mean values from 3 independent experiments. Error bars represent s.e.m. ns=non-significant, *p>0.05, **p>0.01 and *** p>0.001 using multiple comparison with ordinary Two-Ways ANOVA. **c** immunoblot shows level of BRCA1, CtIP and tubulin proteins of irradiated (5 Gy) and non-irradiated U2OS cells after 48 hours of depletion of the correspond genes. **d** U2OS cells in presence of siRNAs against the showed genes upon irradiation (5 Gy) at different time points were blotted for CtIP, RAD52 and α-tubulin.

**Suppl. Fig. 2.**
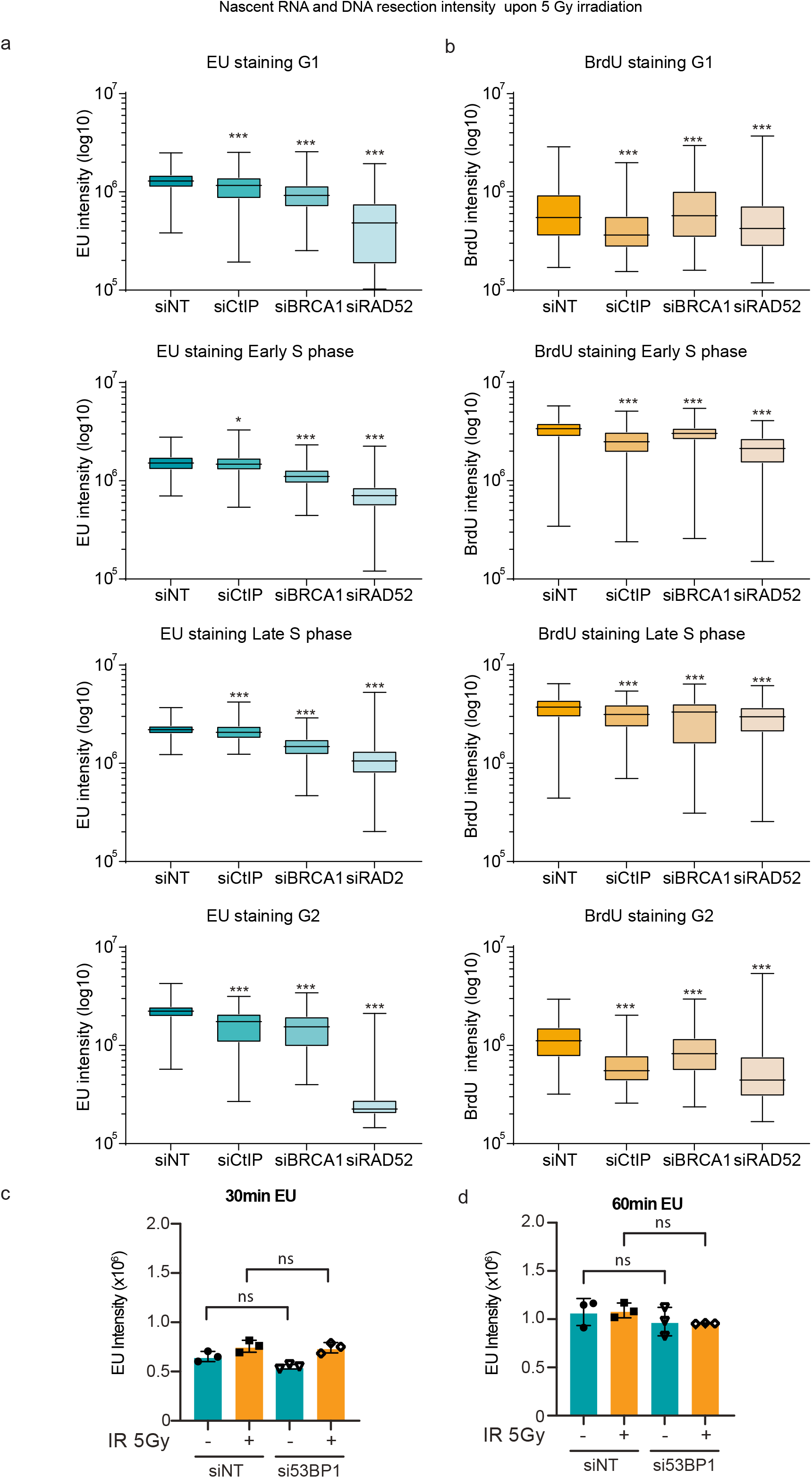
**a** Graph shows nucleoplasm EU intensity in different cell cycle phases in irradiated-U2OS cells. EU component was added upon DNA damage using 5Gy and labelled for 30 minutes. **b** Summary of BrdU intensity in cell cycle phases in U2OS cells treated with BrdU during 24h at the same condition that (a) and BrdU labeling to measure intensity. a-b Showing mean values and 95-5 percentile bars from at least 500 cells. ns=non-significant, *p>0.05, **p>0.01 and *** p>0.001 using multiple comparison with ordinary One-Way. **c-d** Histogram shows nucleoplasm EU intensity in non- and irradiated-U2OS cells with siNT and siRNA against 53BP1. EU component was added upon DNA damage using 5Gy in irradiated cells and labelled for 30 (c) and 60 minutes (d) in both cellular conditions. Showing mean values from 3 independent experiments. Error bars represent s.e.m. ns=non-significant using multiple comparison with ordinary Two-Ways ANOVA.

**Supp. Fig. 3.**
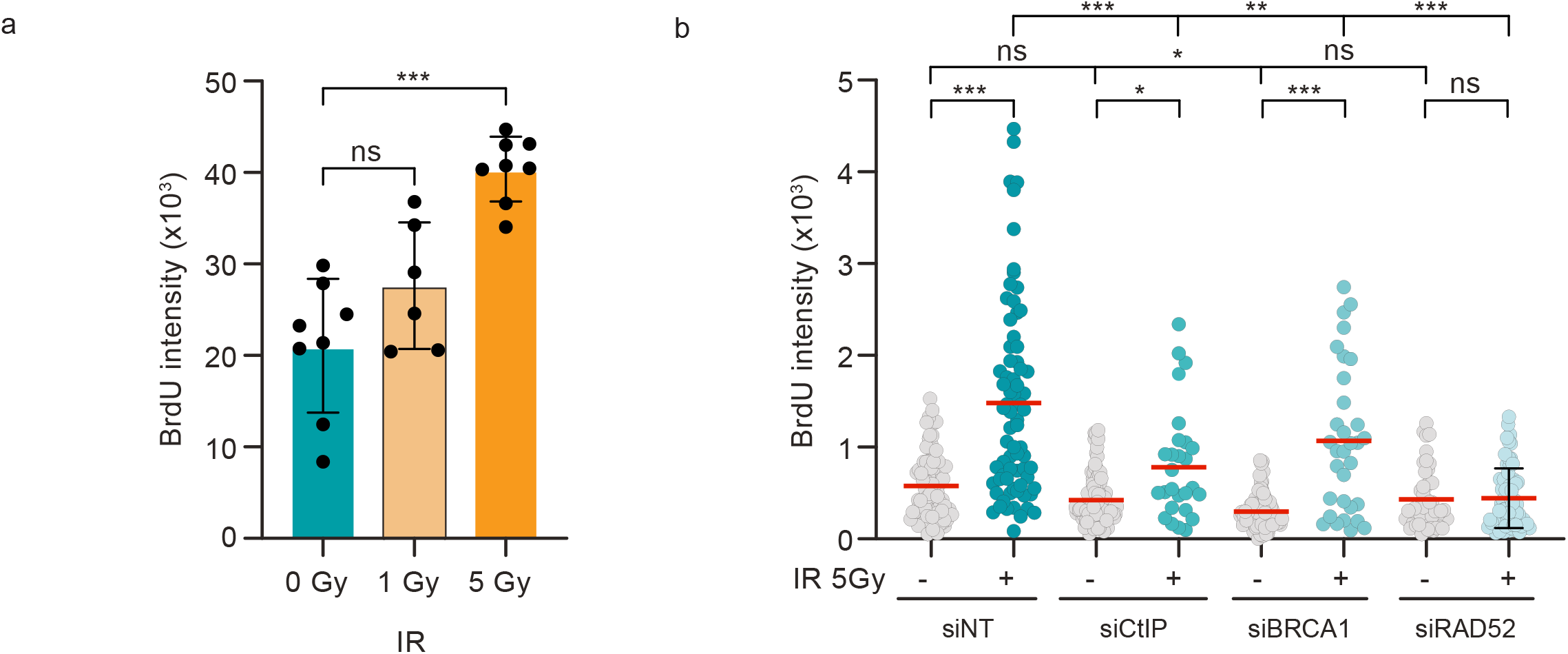
**a** Graph shows average of BrdU intensity from R-SMART samples. Measuring was taken from 0, 1 and 5 Gy samples previously incubated with BrdU during 24 hours. Showing mean values from 3 independent experiments. Error bars represent s.e.m. ns=non-significant using multiple comparison with ordinary One-Way ANOVA. **b** Plot showing BrdU intensity of samples from Hela cells treated with siRNA against CtIP, BRCA1 and RAD52. Measuring was taken from 0 and 5 Gy samples previously incubated with BrdU during 24 hours. At least 25 fields from 3 independent experiments were quantified. ns=non significant, *p>0.05, **p>0.01 and *** p>0.001 using multiple comparison ordinary One-Way ANOVA test.

**Supp. Fig. 4.**
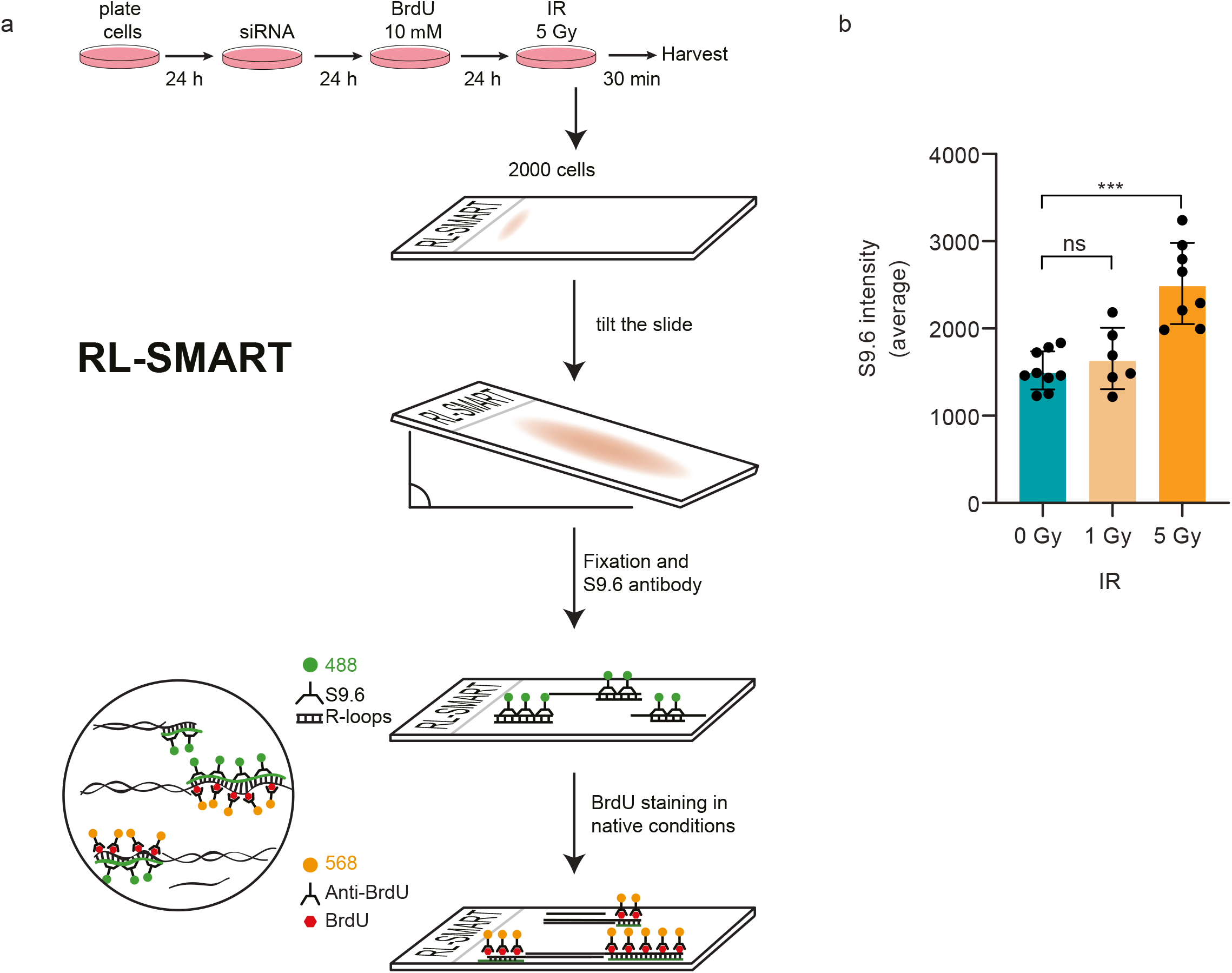
**a** Representative scheme of RL-SMART technology. **b** Graph shows average of S9.6 intensity from RL-SMART samples irradiated with 0, 1 and 5 Gy. Showing mean values from more than 5 samples of 3 independent experiments. Error bars represent s.e.m. ns=non-significant using multiple comparison with ordinary One-Way ANOVA.

**Supp. Fig. 5.**
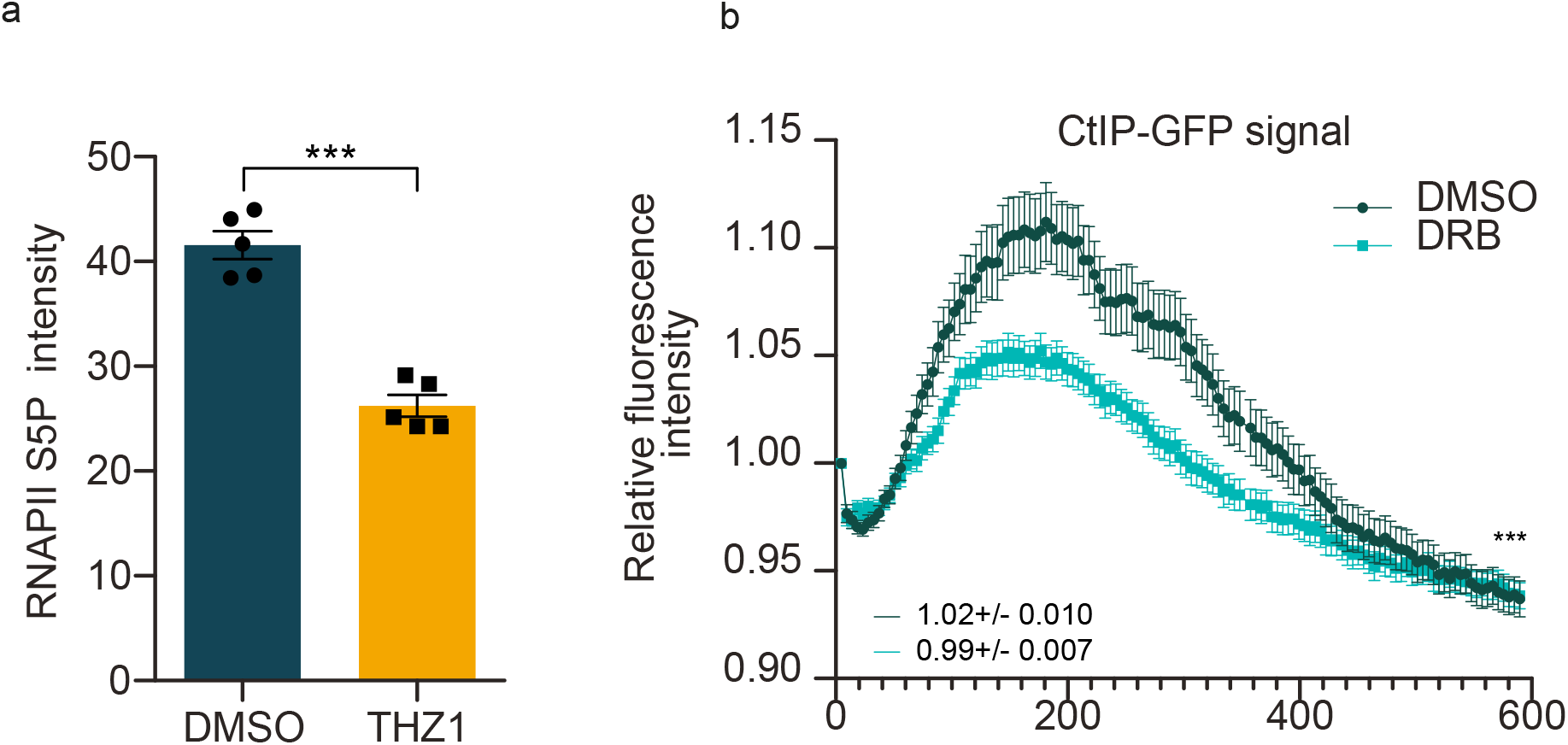
**a** Graph showing intensity of S5P-RNAPII after 2 hours of THZ1 treatment as RNAPII inhibition test. Showing mean values from 5 independent experiments. Showing mean values from 5 independent experiments. Error bars represent s.e.m. ns=non-significant and and *** p>0.001 using ordinary One-Way ANOVA test. **b** Effect of DRB treatment on the kinetics of CtIP recruitment to DNA damage by measuring relative fluorescence intensity of CtIP-GFP during 600 seconds post microlaser irradiation in U2OS cells. The analysis represents the average on n=40 (Ctrl) and n=32 (DRB) nuclei from 3 biologically independent experiments. *** p>0.001 using paired t test.

**Fig. Supp.6.**
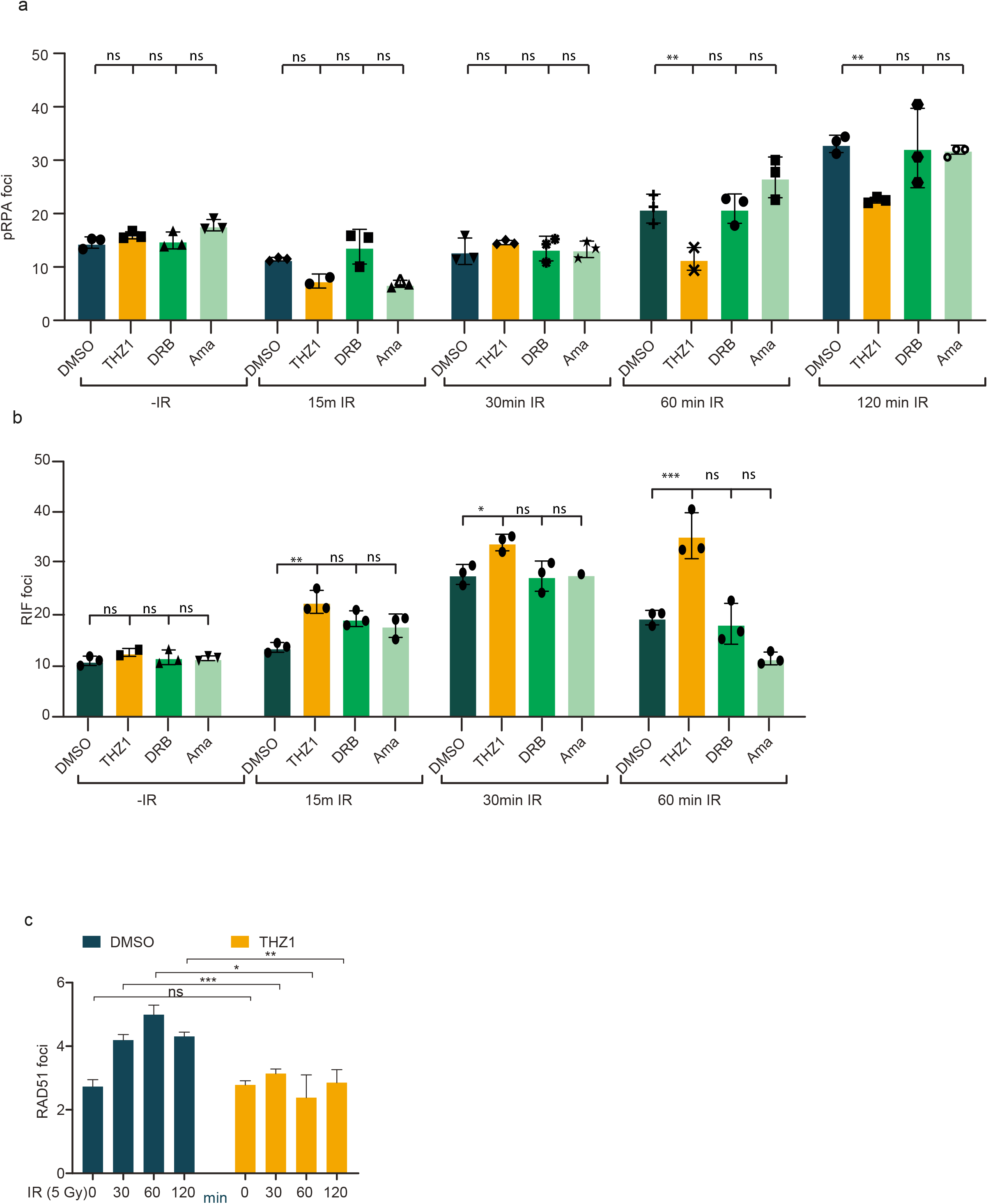
**a** Graph showing S2/S4 phosphorilation of RPA foc upon 5 Gy-irradiation in U2OS cells treated with DMSO, THZ1 (1 µM), DRB and α-amanitin. Bar graph for pRPA foci quantification at 0, 30, 60 and 120 minutes after irradiation. Showing mean values from 3 independent experiment **b** Graph showing RIF1 foci upon 5 Gy-irradiation in U2OS cells treated with DMSO, THZ1 (1 µM), DRB and α-amanitin. Bar graph for RIF1 foci quantification at 0, 30 and 60 minutes after irradiation. Showing mean values from 3 independent experiments At least 2000 cells of ≥ 2 independent experiments were quantified. ns=non-significance *p>0.05, **p>0.01 and *** p>0.001 using multiple comparison with ordinary One-Way ANOVA. **c** U2OS cells treated with THZ1 (1 uM) upon irradiation (5 Gy) at different time points were analysed for RAD51 foci. At least 2000 cells of ≥ 2 independent experiments were quantified. ns=non-significance *p>0.05, **p>0.01 and *** p>0.001 using multiple comparison with ordinary One-Way ANOVA. **e** 53BP1-GFP kinetics upon RNA polymerase II and III inhibitors during 600 seconds post microlaser irradiation in U2OS cells. Graph shows 53BP1-GFP intensity relativized to 1. Cells were treated with RNAPII inhibitor THZ1, 1 µM during 2 hours, and RNAPIII inhibitor during 4 hours prior microlaser irradiation. **f** Same than **e** measuring CtIP-GFP recruitment to DNA damage after RNA polymerase inhibitors.

**Fig. Supp.7.**
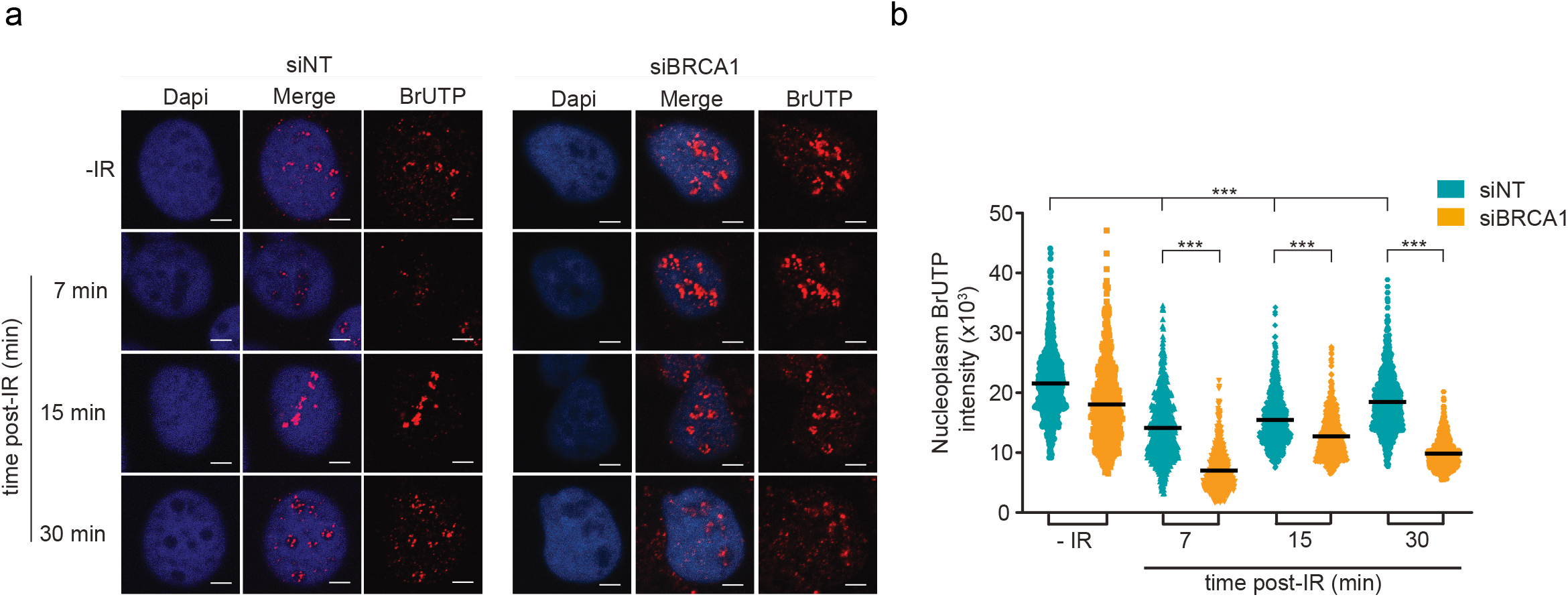
**a** Representative images of *de novo* transcription: BrUTP incorporation into nascent RNA at 7, 15 and 30 min post IR (5Gy) in BRCA1-depleted U2OS cells. Scale bar: 10μm. **b** Dot plot shows nucleoplasm quantification of BrUTP incorporation under the experimental conditions cued in (**a)**. At least 200 cells from each of ≥ 2 independent experiments were quantified. ns=non-significance *p>0.05, **p>0.01 and *** p>0.001 using multiple comparison with ordinary One-Way ANOVA.

